# Simple baselines rival protein language models in dense combinatorial protein landscapes

**DOI:** 10.64898/2026.05.01.722313

**Authors:** Itay Talpir, Sarel J. Fleishman

## Abstract

Computational protein design demands generally applicable models that reliably predict or generate new variants with superior functional properties. A central test of this capability is whether models can extrapolate beyond the mutational regimes represented in their training data. Recent benchmarks have begun to address this question, but the performance of protein language models (pLM) remains less well characterized in densely sampled combinatorial landscapes of protein function that are commonly explored in protein engineering and design. Here, we evaluate zero-shot scoring and supervised prediction across mutational orders, comparing widely used pLMs with simple mutation-identity and evolutionary baselines. We find no consistent advantage associated with pLM architecture or scale, and that pLMs do not consistently outperform conventional baselines. We examined pLM representations and found that much of their variation could be recovered from mutation-identity encoding alone, helping explain the strong performance of such encodings without pretraining. We further found little correspondence between pLM sensitivity to sequence background and experimentally measured functional epistasis. In combinatorial library selection, pLMs likewise showed no consistent advantage over evolutionary scoring. Together, these results indicate that greater pLM complexity does not necessarily improve prediction or experimental prioritization in combinatorial protein design.

## Main

Based on the success and versatility of large language models trained on human text^1^, there has been an explosion of interest in applying similar approaches to proteins. Protein language models (pLMs) typically use Transformer architectures^2–4^, together with pretraining objectives such as masked-token prediction, as in BERT-like models^5^, or autoregressive prediction, as in GPT-like models^6^. In pLMs, self-attention captures relationships between positions in the amino acid sequence, enabling amino acid identities to be predicted from sequence context. These models are typically pretrained on hundreds of millions of naturally occurring protein sequences^3,7^, encoding statistical patterns associated with evolutionary and structural constraints. Thanks in part to their versatility and speed, these models have emerged as efficient and reliable tools for a variety of important bioinformatics tasks^4^, including structure prediction^8–12^, function annotation^7,9,10^, and fitness prediction of mutational variants^13–16^.

In addition to these uses, excitement has surrounded the application of pLMs to design novel protein functions^9,17–19^ or proteins with optimized activities^20–26^. There is, however, a fundamental difference between the above bioinformatics tasks and those of protein design that is likely to limit the applicability of statistical learning to the latter. In many of the former tasks, the challenge is to predict whether a sequence satisfies constraints already common in natural proteins, such as foldability and maintaining some level of native-like activity^27,28^. Thus, a substitution may be assessed by whether it satisfies a loosely defined criterion related to evolutionary plausibility, and a sequence by whether it is likely to maintain a known fold^27^. Prediction, in this sense, is likely to rely mainly on in-distribution interpolation over patterns that are observed in the pretraining data.

By contrast, protein design typically requires generating or predicting the properties of sequences that exhibit new or improved functions relative to those observed in nature^29,30^. The primary challenge is that pLMs are trained on mutations accumulated in natural evolution. Yet, most naturally occurring mutations are neutral rather than functionally advantageous. They may therefore inform design of a space of tolerated mutations but rarely single out advantageous ones, particularly if the design objective is usefulness for a property that is beneficial in artificial contexts. Furthermore, design of improved variants typically demands out-of-distribution (OOD) extrapolation to combinations of mutations that have not been sampled in evolution^30–32^. For instance, successful AI- and physics-based design methods typically introduce dozens of simultaneous mutations to improve protein stability^33–37^, and recent *de novo* enzyme designs have demanded over 100 mutations from any natural protein^38^ and even sequences that lack significant homology to natural proteins^39,40^. In such cases, the pLM faces an OOD prediction problem: it must predict new mutants that are distant from the training data. Although language models can improve OOD prediction robustness in natural language processing^41^, protein design poses a distinct extrapolation challenge: predicting high-order mutants far from experimentally characterized variants. Performance is therefore expected to depend strongly on mutation coverage in the training data and to deteriorate with increasing distance from ground-truth observations^42^. Indeed, a recent analysis found that zero-shot methods, including pLMs, often fail to predict strongly epistatic multi-mutant effects^43^.

pLMs have attracted considerable attention as potentially general-purpose tools for protein design^4,44^, and recent studies have demonstrated their utility in guiding experiments in workflows that combine pretrained models with assay-specific measurements, iterative learning, or explicit modeling of mutational interactions^9,17,22,23,25,45,46^. Recent benchmarks have also begun to evaluate model performance across mutational regimes^31,47–50^. In many protein design and directed-evolution campaigns, the practical goal is to identify variants with improved function, often by recombining a restricted set of candidate mutations. This raises an important question: how much predictive information does pLM pretraining contribute beyond simple mutation-identity and evolutionary baselines in these combinatorial landscapes? The repeated measurement of the same mutations across different sequence backgrounds allows us to evaluate prediction at precisely defined training and test mutational orders and directly examine functional epistasis, a major constraint on protein design^51,52^. Given the substantial resources required to train large models^53–55^ it is important to determine whether their added complexity translates into a consistent advantage for identifying useful, gain-of-function variants.

Here, we evaluate both zero-shot scoring^14,43^ and supervised transfer learning^47,56,57^ on experimentally measured, mutation-dense combinatorial fitness landscapes containing high-order multi-mutant and gain-of-function variants. The datasets span combinatorial libraries in enzymes and fluorescent proteins, and other densely sampled experimental landscapes. We benchmark sequence-based pLMs against simple evolutionary and mutation-identity baselines and additionally evaluate a structure-aware pLM^58^ in compatible analyses. We also test whether pLM representations contain information beyond mutation identity and whether pLM scoring improves model-guided selection of combinatorial sub-libraries. Across these analyses, pLMs show no consistent advantage over simple baselines, and increasing model scale does not consistently improve predictive performance. Much of the mutation-induced variation in pLM embeddings is recoverable from mutation identity, and pLM-nominated sub-libraries do not consistently outperform those nominated by a simple evolutionary baseline.

## Results

### Mutation-dense combinatorial function landscapes

Existing protein fitness benchmarks vary considerably in how extensively they sample combinatorial sequence space, and comparisons with conventional baselines have not always been systematic. ProteinGym, for example, comprises more than 250 experimental assays and over two million experimentally annotated sequences^59^, but most measurements derive from deep mutational-scanning experiments dominated by single and double mutants (Fig. 1a). More recent benchmarks, including FLIP and FLIP2, have expanded these evaluations to higher-order variants and multiple forms of distribution shift^49,50^.

**Figure 1.**
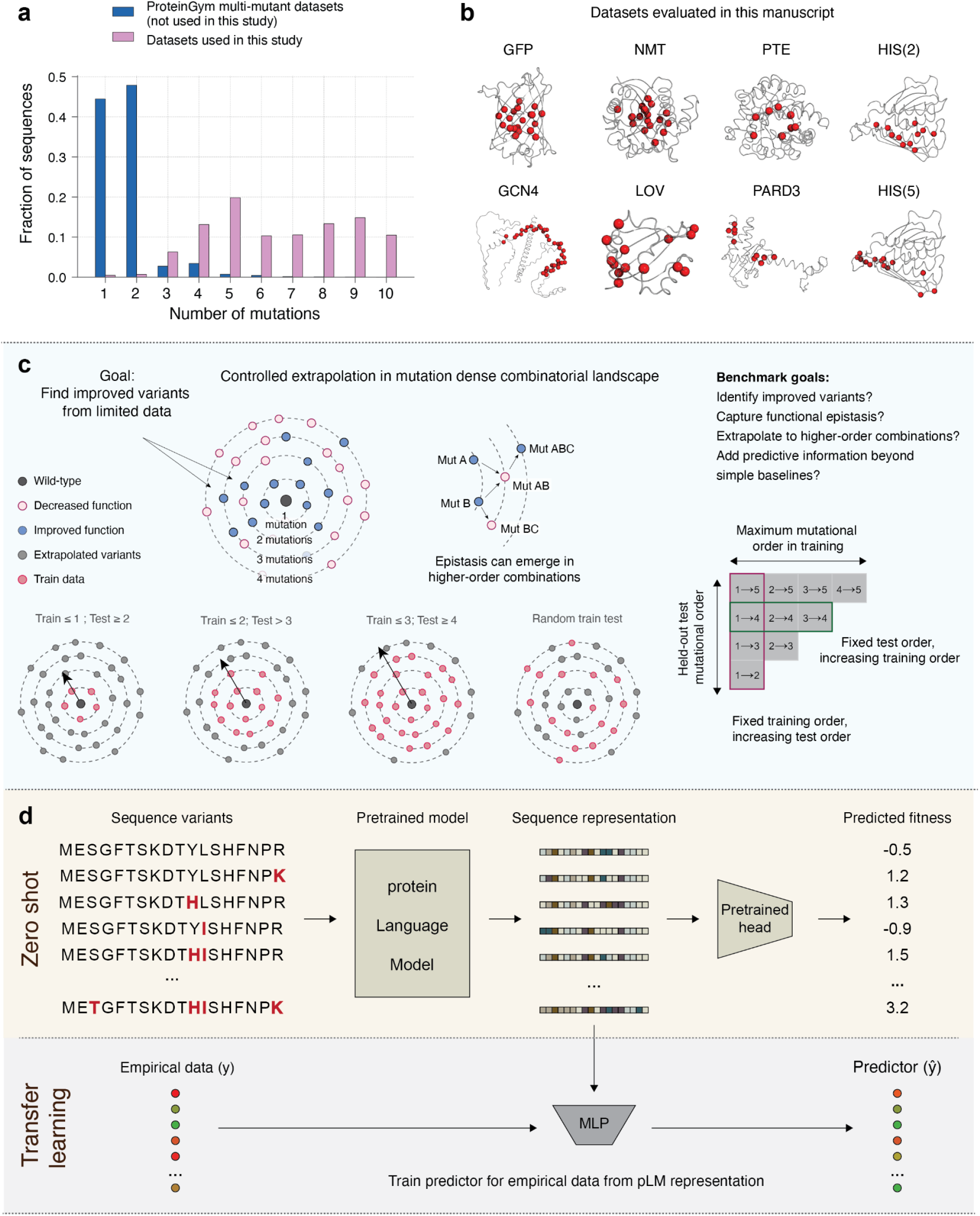
Extrapolation-controlled benchmark for pLMs on mutation-dense combinatorial fitness landscapes. **(a)** Mutational-distance distribution in multi-mutant ProteinGym compared with the datasets used in this study, highlighting limited high-order coverage in standard benchmarks compared to our study. **(b)** Dataset overview (HIS(5), PARD3, GCN4, LOV, HIS(2), PTE, NMT, GFP). HIS(2), HIS(5), PARD3, GCN4 and LOV were obtained from ProteinGym; HIS(2) and HIS(5) represent distinct combinatorial regions of the same assay. GFP and PTE are publicly available. Dataset sources, assay-specific phenotypes, availability, and analysis usage are summarized in Supplementary Table 1. **(c)** Controlled extrapolation framework for mutation-dense combinatorial landscapes. Models are trained on lower-order variants and evaluated on held-out higher-order combinations, enabling systematic tests of improved-variant prediction, epistasis, and extrapolation with increasing mutational order. Bottom, representative controlled train-test regimes and a random-split comparison; right, the mutational-order train-test combinations evaluated. **(d)** Two common pLM use-modes in protein design. Top: zero-shot scoring of variants with a pretrained model (without assay-specific data). Bottom: transfer learning, where pLM representations are used to train a supervised predictor on empirical measurements.

In protein design and machine-learning-assisted directed evolution, candidate mutations, often at binding or catalytic sites, are frequently recombined to explore restricted sequence spaces. Despite their practical relevance, densely sampled combinatorial landscapes of function are usually not analyzed. The recent FLIP and FLIP2 benchmarks include notable combinatorial examples, such as GB1 and TrpB, but these primarily involve combinations across three or four positions at a time. Here, we focus on functional landscapes spanning larger sets of selected positions, with substitutions repeatedly sampled across diverse sequence backgrounds (Supplementary Fig. 1c,d). This sampling structure enables controlled extrapolation across mutational orders and direct analysis of functional epistasis. It also allows us to investigate whether pLMs provide predictive information beyond mutation-identity and evolutionary baselines, and whether they improve combinatorial library selection (Fig. 1b,c).

Three of the landscapes we analyzed were generated using the FuncLib^37,60,61^ and htFuncLib^33^ protein-design frameworks, which combine evolutionary information with Rosetta-based atomistic calculations to generate multi-mutant variants predicted to fold stably while maintaining key residues in functionally competent orientations. Rather than targeting a specific functional outcome, these methods generate panels of functionally related variants and have yielded substantial improvements in enzyme selectivity, catalytic rates, and specificity^62–64^, including in *de novo* enzyme design^38^. The resulting landscapes contain higher-order combinations across 8 to 22 designed positions. The htFuncLib GFP landscape is especially dense, containing thousands of experimentally validated combinatorial variants, including mutants with up to eight substitutions. Many variants exhibit strong sign epistasis^37,51^, in which mutation effects can change or even reverse depending on the sequence background^43^. Such interactions are critical in protein engineering as they can constrain the emergence of improved functions during protein design and natural or laboratory evolution^51,52,65^. We identified epistatic mutation pairs by comparing their observed effects to those expected from lower-order variants (Supplementary Fig. 5).

To broaden the benchmark beyond FuncLib-derived landscapes, we also analyzed four ProteinGym datasets: LOV^66^, PARD3^67^, GCN4^68^, and HIS^69^ (Fig. 1b). These were selected for their dense coverage of multi-mutant combinations. The HIS assay contains distinct combinatorial regions, of which we analyzed the second and fifth regions, labeled as HIS(2) and HIS(5). Together, the landscapes span 10 to 38 designed positions, range from tens to thousands of measured sequences, and encompass diverse proteins and experimental phenotypes. We also included the AAV capsid landscape from the original FLIP benchmark in selected analyses, depending on the mutational coverage required (Supplementary Table 1). The combinatorial coverage, higher-order variants, and gain-of-function signal in our benchmark enable matched comparisons between pLMs and simple baselines. We evaluate two common modes of pLM application, zero-shot scoring and supervised transfer learning (Fig. 1d), with particular emphasis on prediction and extrapolation across mutational orders.

### Evolutionary scoring rivals pLMs in zero-shot predictions

Previous studies reported that pLMs provide useful zero-shot rankings of protein variants^14,25^. We therefore evaluated this claim in our combinatorial landscapes using two complementary scoring procedures across eight pLMs^10,11,14,22,23^ (Fig. 2a, Supplementary Fig. 2-3, Supplementary Table 2). In the first procedure, fixed-context additive scoring, all varied positions were masked in a common sequence context. The sequence was then passed through the pLM, and mutation-specific log-likelihood differences for the substitutions in each variant were summed. In the contextual, background-dependent procedure, only positions mutated in the evaluated sequence were masked, such that the conditioning context changed with the mutational background (see Methods). For ProGen2, the corresponding contextual score was computed using left-to-right decoding, allowing later positions to be scored conditional on preceding ones. This second procedure allows substitution scores to depend on sequence background and tests whether background-dependent scoring improves prediction relative to the fixed-context additive formulation.

**Figure 2.**
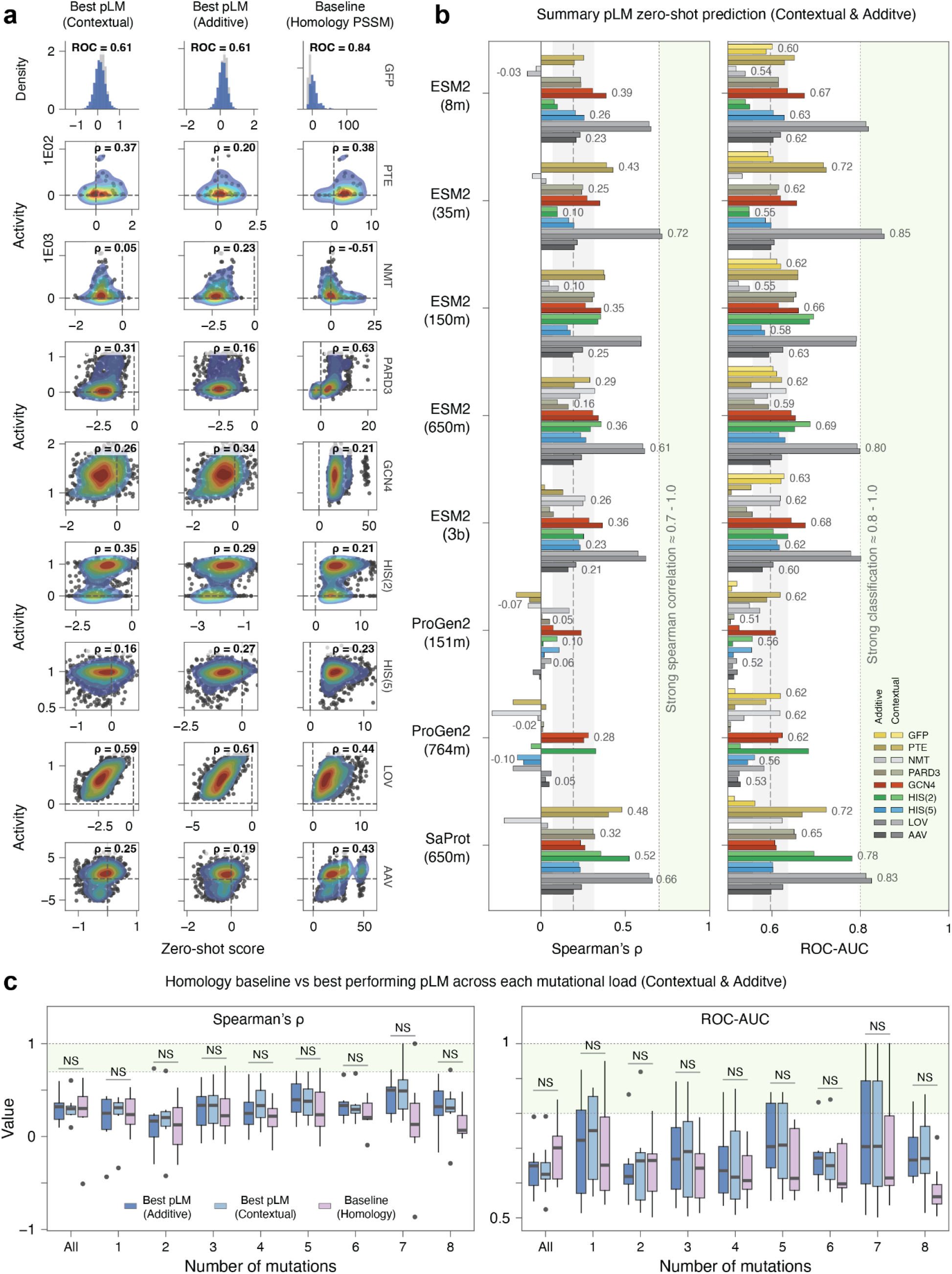
PSSM baseline matches zero-shot pLM scores in mutation-dense combinatorial landscapes. **(a)** Representative zero-shot comparisons of the best-performing pLM scores and the PSSM baseline across the evaluated landscapes. For GFP, distributions of predicted scores for functional and nonfunctional variants are evaluated by ROC AUC. For the remaining datasets, scatter plots show experimental phenotypes versus zero-shot scores, with Spearman ρ reported for each comparison **(b)** Cross-dataset summary of zero-shot performance on mutation-dense multi-mutant landscapes from FuncLib and ProteinGym. Left, Spearman correlation with quantitative phenotypes; right, ROC AUC for classification. Gray shading indicates the mean ± s.d. across models and datasets. Dataset identity is indicated by the bar color (see legend). **(c)** pLM zero-shot performance compared with a PSSM baseline, aggregated across datasets for ROC AUC and Spearman ρ. Differences are not significant (Kruskal–Wallis test; NS, P > 0.05).

Across the mutation-dense landscapes, additive and contextual scoring produced qualitatively similar results. Classification performance was modest, with an average ROC AUC approximately 0.6. Correlations with quantitative phenotypes were low, with an average Spearman *ρ* approximately 0.2 (Fig 2b). Neither scoring procedure showed a systematic advantage over the other. We next asked whether pLM zero-shot scores outperformed established evolutionary sequence priors by comparing them to position-specific scoring matrices (PSSMs). This comparison is particularly relevant because evolutionary sequence models have been used successfully in protein design and engineering for decades^70^ and effective models for design can be constructed even for proteins with several dozen diverse natural homologs^36,71^. To compare the pLMs favorably, for each dataset and mutational order we selected the best-performing pLM under each zero-shot scoring procedure. Still, pLMs did not significantly outperform PSSMs in either ROC AUC or correlation coefficient (Fig. 2c). The same result was obtained using an additional additive zero-shot procedure, in which each mutated position was scored individually and the resulting mutation-level scores were summed across the variant. We concluded that in these mutation-dense combinatorial landscapes, conventional evolutionary baselines are competitive with pLM zero-shot scoring.

### pLM sensitivity to sequence background poorly reflects functional epistasis

We next asked whether the sequence-background sensitivity of zero-shot scoring by pLMs reflects experimentally measured functional epistasis. We compared contextual scores, obtained by jointly masking all mutated positions, with additive scores, obtained by masking and scoring each mutation independently in the wild-type background. We correlated the signed difference between these scores with experimental pairwise epistasis (ε_AB), defined as the deviation of the double-mutant phenotype from the additive expectation based on its constituent single mutants. Correlations were generally weak across datasets and pLMs, with an average Spearman *ρ* close to zero (Supplementary Fig. 7a-b).

To avoid assumptions about the numerical scale on which mutational effects combine, we also examined unexpected functional outcomes, to which we refer as qualitative epistasis. We identified mutation pairs whose binary phenotype (active/inactive) deviated from an AND-rule expectation based on their constituent single mutants (Methods). We further identified higher-order variants in which additional mutations reversed the measured double-mutant phenotype (Supplementary Fig. 5). The prevalence of these interactions and the diversity of reversing backgrounds varied across datasets. LOV exhibited fewer such pairs and simpler reversal patterns, consistent with its greater predictability by linear models (Supplementary Fig. 6).

We then tested whether the magnitude of contextual-minus-additive score differences distinguished qualitatively epistatic pairs, and whether contextual scoring improved phenotype prediction in variants containing these interactions. Discrimination was generally near chance, and contextual scoring did not consistently improve phenotype correlations (Supplementary Fig. 8-9). These results indicate that the sensitivity of pLM zero-shot scores to sequence background does not reliably reflect experimentally measured functional epistasis. These findings are consistent with a previous report^43^ that demonstrated reduced zero-shot prediction accuracy on experimentally epistatic variants.

### Training data, not pLM type, determines performance

We then tested whether pLM representations improve supervised transfer learning^47,56,57^ in settings that simulate combinatorial protein design. Here, models are trained on experimentally measured low-order variants and evaluated on previously unseen high-order combinations. For each pLM, we computed sequence embeddings and trained a supervised predictor on the measured phenotypes (Fig. 1d bottom). Because transfer-learning performance may depend on the downstream learning algorithm, we compared ridge regression, XGBoost, and multilayer perceptron (MLP) predictors. MLPs generally performed best across the evaluated datasets and metrics and were therefore used for subsequent supervised benchmarks (Supplementary Fig. 11; Methods). We also restricted pLM representations to experimentally varied positions rather than the entire sequence. This improved performance relative to full-sequence embeddings and reproduced the supervised performance reported in the original LOV study^66^.

The dense combinatorial structure of the landscapes in our benchmark allows controlled separation of training and test variants by mutational order (Fig. 1c). We evaluated eight pLMs spanning different architectures, parameter counts, and training objectives in two complementary settings. First, we fixed the mutational order of the held-out test set and varied the amount of lower-order information available for training. For example, models could be evaluated on the same set of quadruple mutants after being trained on progressively larger subsets of lower-order variants (Fig. 3a,b; Supplementary Fig. 12a,c). Some of the metrics we used are directly relevant to protein design. In particular, Precision@100 (Fig. 3b) simulates a practical design setting in which, given experimental budget limitations, the model nominates the 100 variants most likely to be functional. Second, we fixed the lower-order training data and evaluated each model on progressively higher mutational orders. For example, models trained on single and double mutants could be compared across triple, quadruple, and higher-order test variants (Supplementary Fig. 12b,d). Both settings represent extrapolation prediction scenarios^31^.

**Figure 3.**
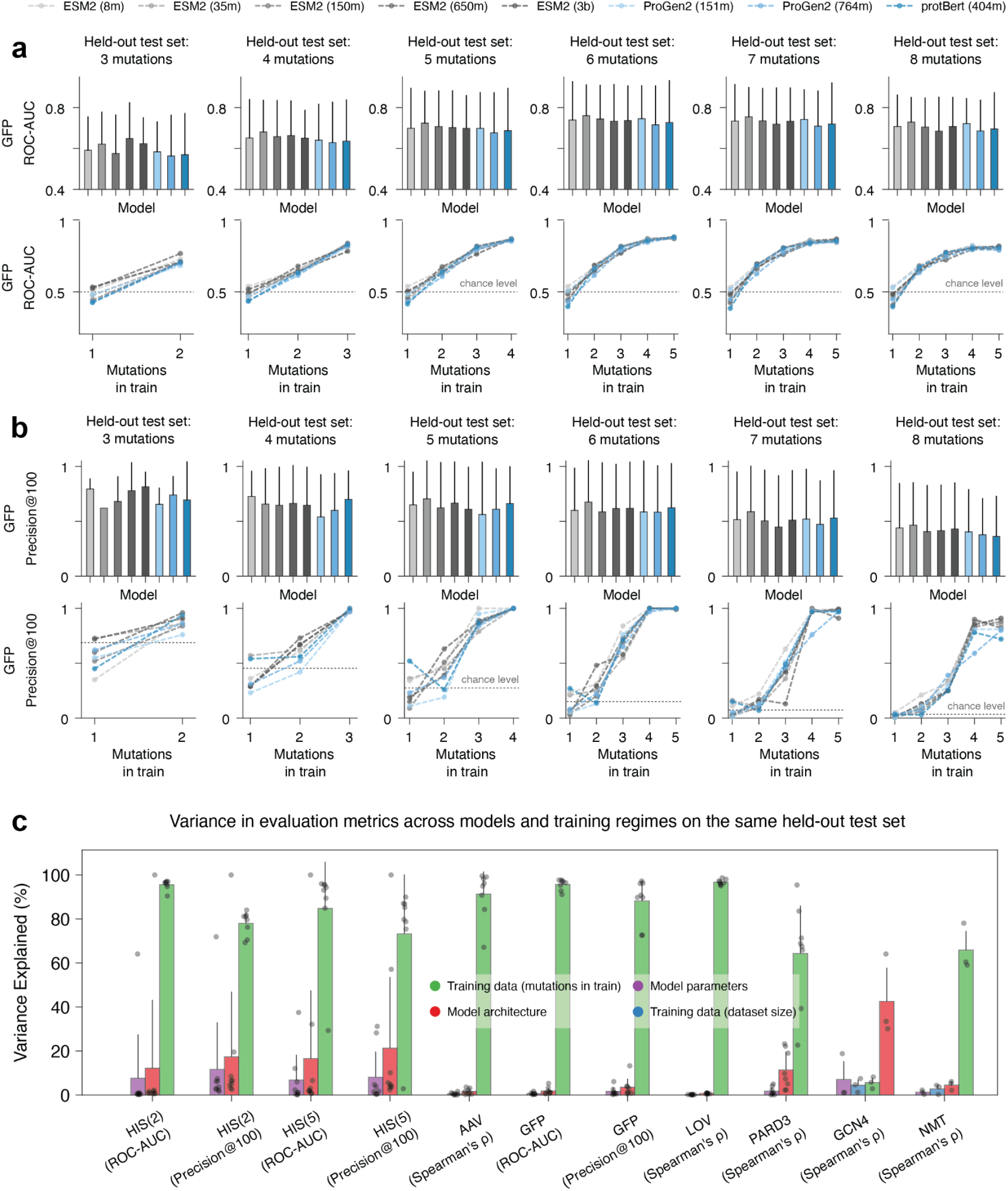
Under controlled extrapolation, pLMs are largely indistinguishable and performance is dominated by the train–test mutational gap. **(a)** ROC AUC for the GFP dataset under controlled extrapolation, evaluated on held-out test sets with 3–8 mutations. Each panel corresponds to a fixed test mutational order (e.g., all triple mutants). For a given test order, each pLM boxplot aggregates performance across all training regimes evaluated at that test order (for example, training on 1-mutants only, 1+2, 1+2+3, etc., and testing on 4-mutants). Lines summarize performance as a function of the maximum mutational order included in training. The top and bottom subpanels show the same underlying splits and results, displayed with complementary groupings: once as a function of training mutational order and once as a function of pLM identity. **(b)** Same as **(a)**, evaluated using Precision@100. Chance level is shown by a horizontal dashed line **(c)** Variance explained analysis across models and training regimes for matched held-out sets, showing that training regime (mutational order/distance coverage) explains most variance relative to model scale, architecture, and training budget.

GFP illustrates why this extrapolation problem can be particularly challenging. Nearly all measured single mutants are functional, whereas combinations of individually tolerated mutations are often non functional because of negative sign epistasis (Supplementary Fig. 5a, Supplementary Fig. 10). Consequently, models trained only on single mutants receive no direct information about which higher-order combinations will fail, and their ROC AUC can remain close to random expectation. In this setting, pretrained representations could, in principle, provide information about mutational compatibility that is absent from the measured lower-order phenotypes. The controlled benchmark provides a direct test of whether such information improves prediction of unseen combinations.

Across datasets, the performance of pLMs overlapped, with no clear trend indicating that any model outperformed others (Fig. 3a,b, top; Supplementary Fig. 13). Performance was instead strongly dependent on the separation between the mutational orders represented in training and those used for evaluation. We quantified these effects across both controlled-extrapolation settings using analysis of variance. In approximately 93% of model comparisons, differences between pLMs were not statistically significant, and no model family or scale showed a consistent advantage (Supplementary Fig. 13a-c). By contrast, differences between training regimes evaluated on the same held-out test set were statistically insignificant in only approximately 5% of comparisons (Supplementary Fig. 13d), indicating that the composition of the training data had a substantially larger effect on predictive performance than pLM type.

Finally, we used factor-wise variance attribution analysis to identify the main drivers of performance differences (Fig. 3c). Across the landscapes, variation associated with the training mutational regime was substantially larger than variation associated with pLM type, architecture or parameter-count bin. The observation that increasing parameter count did not translate into improved performance is surprising. For example, ESM-8M (8 million parameters) performed similarly to, and often better than, ESM-3B (3 billion), despite a nearly three orders of magnitude difference in parameter count and proportional differences in cost to develop and train^53,54^. These results suggest that scaling to more computationally intensive models does not yield a clear or consistent performance benefit in extrapolative settings of variant prediction, which are central to protein design. More broadly, this is consistent with recent work showing that protein-engineering predictors can be limited less by model architecture than by the amount and diversity of the available training data^42^.

### One-hot baseline rivals pLMs in transfer learning

We next asked whether pLM representations outperform a computationally simple baseline in the same supervised prediction setting. We repeated the controlled-extrapolation analyses described above, comparing the best-performing pLM for each dataset and metric with a one-hot encoding (OHE) of mutation identity followed by the same class of MLP predictor (Fig. 4a). This comparison deliberately favors the pLMs, as a different pretrained representation could be selected for each dataset and metric, whereas OHE simply records the amino-acid identities present at the experimentally varied positions. To summarize performance across train-test regimes, we calculated the ratio between OHE and the selected pLM for each comparison (Fig. 4b). OHE matched or outperformed the selected pLM in nearly all settings, and the aggregate distribution favored OHE in all but one dataset-metric combination.

**Figure 4.**
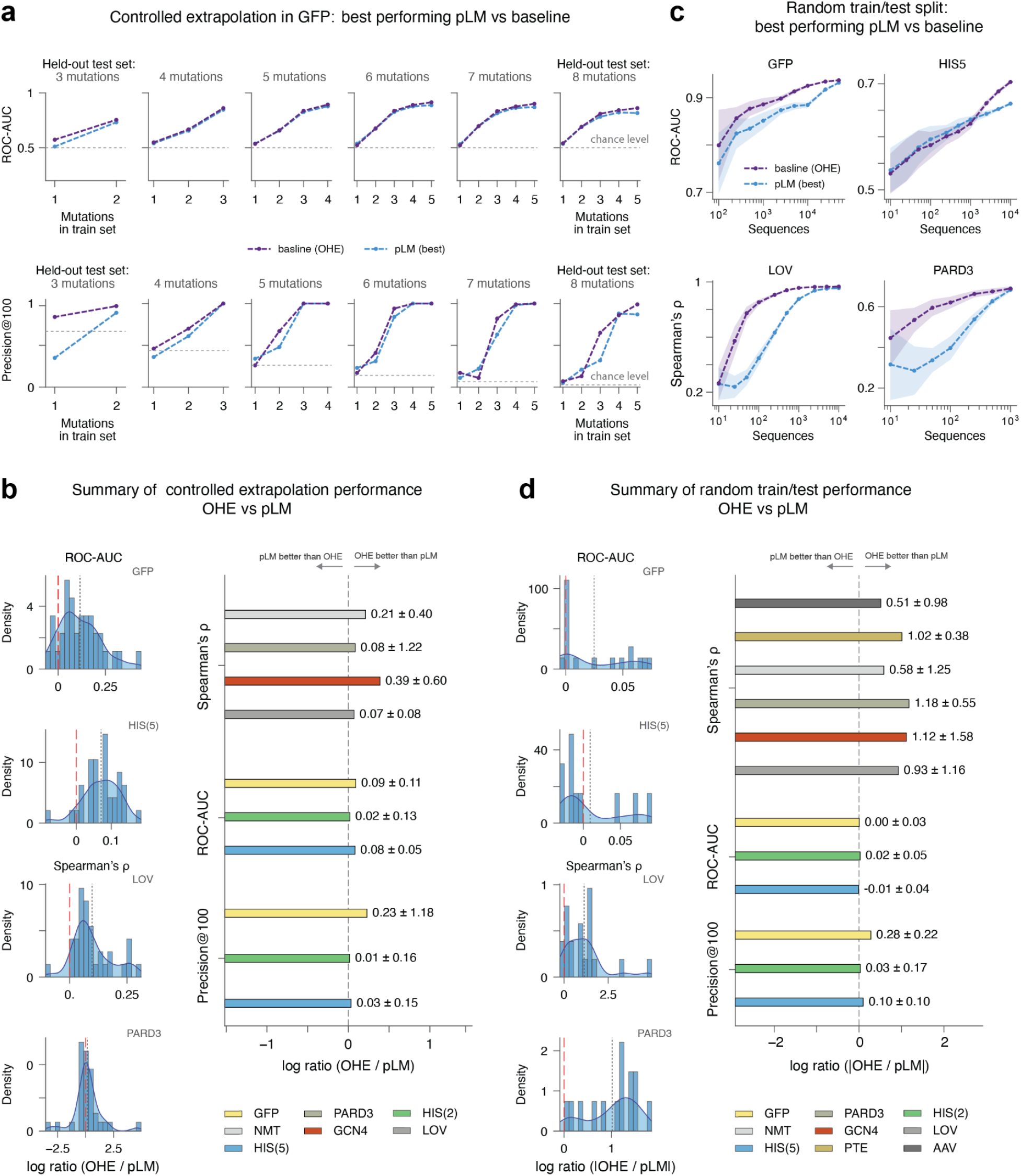
One-hot mutation encoding rivals pLM representations in controlled extrapolation and random train-test splits. **(a)** Top: ROC AUC for the GFP dataset under controlled extrapolation comparing OHE+MLP to the best-performing pLM for each dataset/metric across held-out test mutational orders. OHE matches or exceeds the best pLM across most regimes. Bottom: Same as top, evaluated using Precision@100 under the same controlled-extrapolation evaluation. **(b)** Summary of OHE versus pLM performance across controlled-extrapolation regimes. Left, distributions of the log performance ratio between OHE and the selected best-performing pLM across train-test combinations for representative datasets and metrics. Right, dataset-level summaries across Spearman ρ, ROC AUC, and Precision@100. Positive log ratios indicate better performance by OHE and negative values indicate better performance by the pLM. Bars show the mean ± standard deviation across the corresponding comparisons. **(c)** Comparison of OHE and the best-performing pLM under conventional random train-test splitting as a function of the number of experimentally measured training sequences. Representative classification and regression datasets are shown, with ROC AUC for GFP and HIS(5) and Spearman’s ρ for LOV and PARD3 **(d)** Corresponding summary for random train-test splits across training-set sizes. Left, distributions of OHE-to-pLM performance ratios for representative datasets; right, aggregate dataset-level comparisons for Spearman ρ ROC AUC, and Precision@100. As in the controlled-extrapolation analysis, the best-performing pLM was selected separately for each dataset and metric before comparison with OHE. Despite this pLM-favorable model-selection procedure, OHE remained competitive with or outperformed the selected pLM across most datasets, metrics, and evaluation regimes.

We then evaluated the same representations under conventional random train-test splits, where training and test variants were sampled from the same mutational distribution rather than separated systematically by mutational order. For this analysis, we used the pLM that performed best overall across datasets in the controlled-extrapolation analyses, rather than reselecting a pLM under the random-split setting. OHE again matched or exceeded this pLM across a broad range of training-set sizes (Fig. 4c), and the corresponding aggregate performance ratios favored OHE across nearly all datasets and metrics (Fig. 4d).

Because supervised performance can depend on model selection, we tested four alternative train-validation-test procedures using two representative pLMs. For controlled extrapolation, we compared validation within the low-order training regime, validation on the next mutational order to explicitly select for extrapolation, and a deliberately optimistic post hoc procedure that selected the best architecture using the broader dataset before refitting under the controlled protocol. For random splits, we compared model selection within each subsample with selecting an architecture once using the broader dataset. OHE remained highly competitive and often surpassed pLM representations across these procedures (Supplementary Fig. 14; Methods).

We next tested whether this result depended on how the pLM representation or downstream predictor was constructed. Using representative pLMs, we compared alternative embedding layers, pooling strategies, and supervised architectures with the OHE baseline (Supplementary Fig. 15). The relative performance of OHE and pLM representations was similar across these choices. The fixed final-layer representation used in our supervised benchmarks performed similarly to the best embedding scheme selected separately within each training regime.

Because frozen representations may underestimate the benefits of task-specific adaptation, we also evaluated low-rank adaptation (LoRA) fine-tuning under the same controlled extrapolation protocol. LoRA did not outperform OHE in this setting (Supplementary Fig. 16-17).

Together, these results emphasize that, in protein design tasks on mutation-dense combinatorial landscapes, pLM performance is limited primarily by the training data, and, crucially, that simple baselines rival state-of-the-art pLMs. Recent analyses of pLM performance on low-order mutants are consistent with our findings, indicating that simple baselines, such as OHE, are competitive with, and often surpass, pLM-based approaches across low- and high-order mutants^21,43,47,50,72,73^.

### pLM embeddings substantially overlap with mutation identity

The competitive performance of OHE raises a question that prediction benchmarks alone cannot resolve: how much of the variation in pretrained pLM representations across these landscapes is already explained by mutation identity? OHE does not incorporate pretrained information about protein sequence or evolutionary context; it solely records the presence of specific mutations. We therefore compared pairwise distances in both OHE and pLM representation spaces. Across the landscapes, pairwise Hamming distances in OHE space were moderately to strongly correlated with pairwise cosine distances between pLM embeddings, with Spearman correlations ranging from approximately 0.4 to 0.8 (Fig. 5a,b). Thus, despite dramatic differences in architecture, complexity, and training costs, these representations are, to some extent, similar in how they organize local sequence spaces.

**Figure 5.**
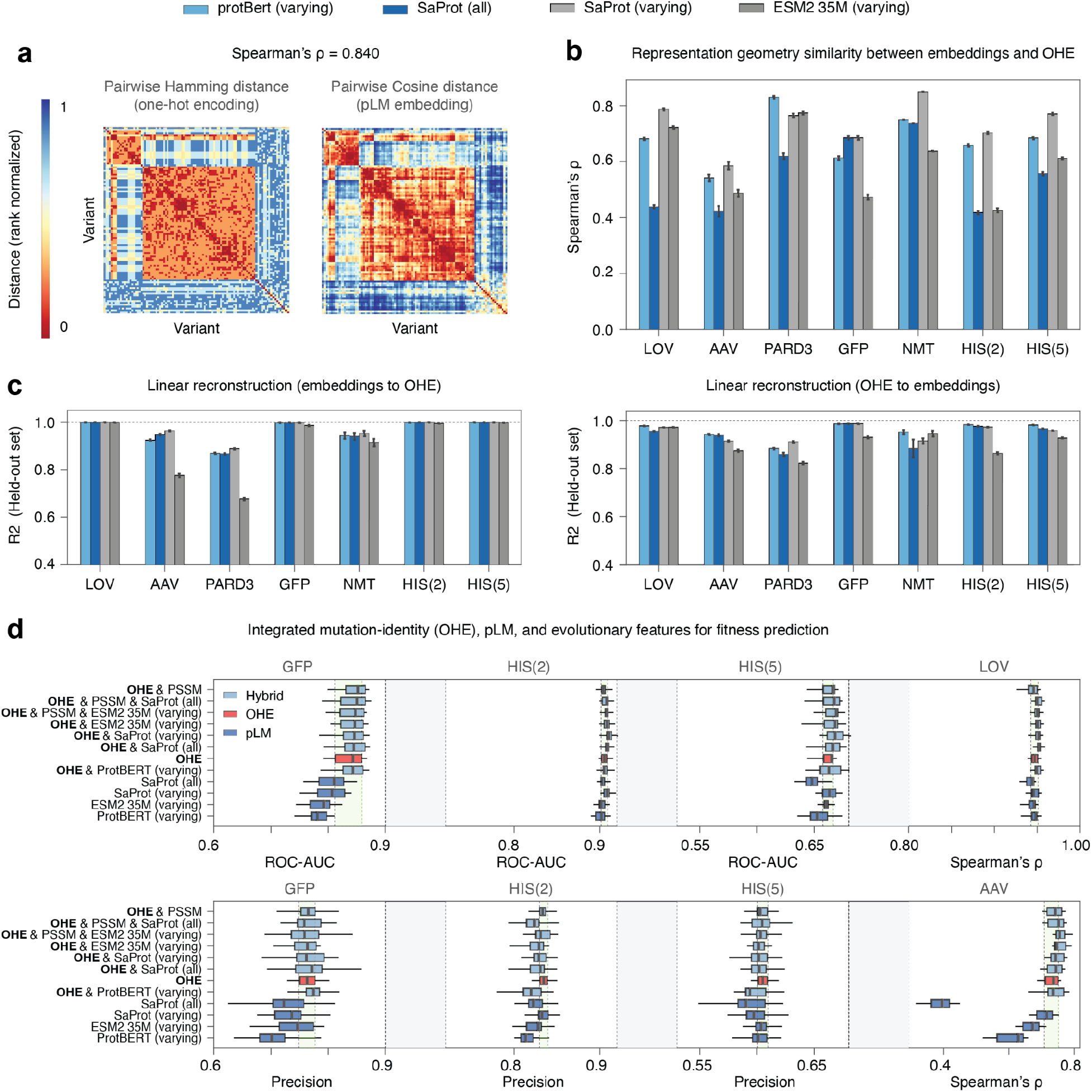
Much of the variation in pLM representations is predictable from mutation identity in dense combinatorial landscapes. **(a)** Representative comparison of pairwise distances between variants encoded by one-hot mutation identity (OHE) and by pLM embeddings (SaProt, PARD3 dataset). Pairwise Hamming distances in OHE space are shown alongside pairwise cosine distances between pLM embeddings, with variants ordered identically in both matrices. Distances are rank-normalized for visualization. The Spearman correlation between the two pairwise-distance matrices is shown above, illustrating the similarity of the sequence-space geometry represented by OHE and the pLM. **(b)** Similarity between OHE and pLM representation geometries across landscapes. For each dataset, Spearman ρ was calculated between pairwise Hamming distances in OHE space and pairwise cosine distances in the indicated pLM representation. Representations include ProtBert, ESM2-35M, and SaProt. "Varying" indicates that embeddings were restricted to experimentally varied positions, whereas "all" indicates representations derived from all sequence positions. Pairwise geometry was substantially correlated across datasets and representation choices. **(c)** Linear reconstruction between pLM and OHE representations. Left, ridge-regression models were trained to reconstruct OHE representations from pLM embeddings and evaluated on held-out variants. Right, the reciprocal analysis reconstructs pLM embeddings from OHE representations. Models were trained on half of the variants and evaluated on the remaining held-out variants. Bars show held-out R^2^ for the indicated pLM representations and datasets. High reconstruction accuracy in both directions indicates that a substantial component of each representation is linearly recoverable from the other. **(d)** Fitness prediction using individual and combined mutation-identity, pLM, and evolutionary representations. OHE, PSSM, and the indicated pLM representations were evaluated individually or concatenated before training the downstream predictor. Results are shown for GFP, HIS(2), HIS(5), LOV, and AAV using the corresponding classification or regression metric. The red box denotes the OHE-only baseline, while the remaining boxes show individual pLM representations or hybrid representations containing OHE together with pLM and/or PSSM features. Adding pLM embeddings to OHE or evolutionary features yielded only small and inconsistent improvements across datasets and evaluation metrics, consistent with substantial redundancy between explicit mutation identity and the pLM representations in these densely sampled combinatorial landscapes.

We next investigated whether the observed overlap was sufficient for one representation to predict the other. For each dataset, we randomly selected variants and trained ridge-regression linear probes on half of them to reconstruct OHE from pLM embeddings and *vice versa*. Reconstruction performance was evaluated on held-out variants. Strikingly, predictability was high in both directions: *r*² values ranged from approximately 0.6 to 0.99 for pLM-to-OHE reconstruction and from 0.8 to 0.99 for OHE-to-pLM reconstruction (Fig. 5c). Thus, much of the variation in pLM representations across these landscapes is linearly recoverable from explicit mutation identity.

Because representational reconstruction does not directly measure predictive usefulness, we next tested whether adding pLM embeddings to mutation-identity or evolutionary features improves phenotype prediction. We similarly performed an extensive model-selection experiment to identify optimal downstream predictors (see Methods). Incorporating pLM embeddings with OHE or evolutionary features resulted in only minor and inconsistent improvements across datasets (Fig. 5d). Collectively, these analyses indicate that, within mutation-dense combinatorial landscapes relevant to protein design, much of the variation in pLM representations is recoverable from explicit mutation identity, while combining these representations with mutation-identity or evolutionary features provides little consistent additional benefit for predicting functional phenotypes.

### Evolutionary scoring rivals pLMs in activity-landscape exploration

Recent studies have suggested that pLMs can guide experimental exploration of protein fitness landscapes^20,21,25^. In this setting, the goal is to focus experimental screening on regions of sequence space that are more likely to contain functional or improved variants^74–76^. We asked whether pLM zero-shot scores are superior relative to a simple evolutionary baseline in two related design tasks: prioritizing individual mutations and defining combinatorial sub-libraries for experimental screening. We extended this analysis to include the TrpB and AlphaAmyl datasets^77,78^ to test its scope.

We first evaluated whether model scores recover empirically favorable mutations. For each mutation, we calculated an empirical mutation score as the ratio of the mean activity of variants containing that mutation with variants in which it was absent (Supplementary Fig. 18a; Methods). We then asked whether pLMs and PSSMs preferentially ranked mutations with high empirical scores and whether model rankings correlated with empirical mutation effect across the full set of mutations. pLMs and PSSMs showed some ability to prioritize favorable mutations, but correlations were generally modest and neither showed a consistent advantage over the other (Supplementary Fig. 19a left). Importantly, in most landscapes both pLMs and PSSMs distinguished mutations associated with the highest and lowest activities (Supplementary Fig. 19a, left and middle), As in our previous analyses, larger pLMs were not consistently better than smaller ones in mutation ranking.

We next simulated a typical protein engineering decision: whether pLMs nominate sets of mutations that define combinatorial screening libraries that comprise more functional variants than those selected by conventional evolutionary scoring. For each empirical landscape, each model nominated a restricted set of mutations across the designed positions, thereby defining a combinatorial sub-library. For each model-nominated sub-library, we considered experimentally measured variants that contained only mutations from the nominated set, with at most one additional mutation (Supplementary Fig. 18c; Methods). Eligible variants were repeatedly sampled at increasing screening budgets. If a model identifies a useful sub-library, variants sampled from that library should show higher activity and contain more gain-of-function sequences (Figure 6b).

**Figure 6.**
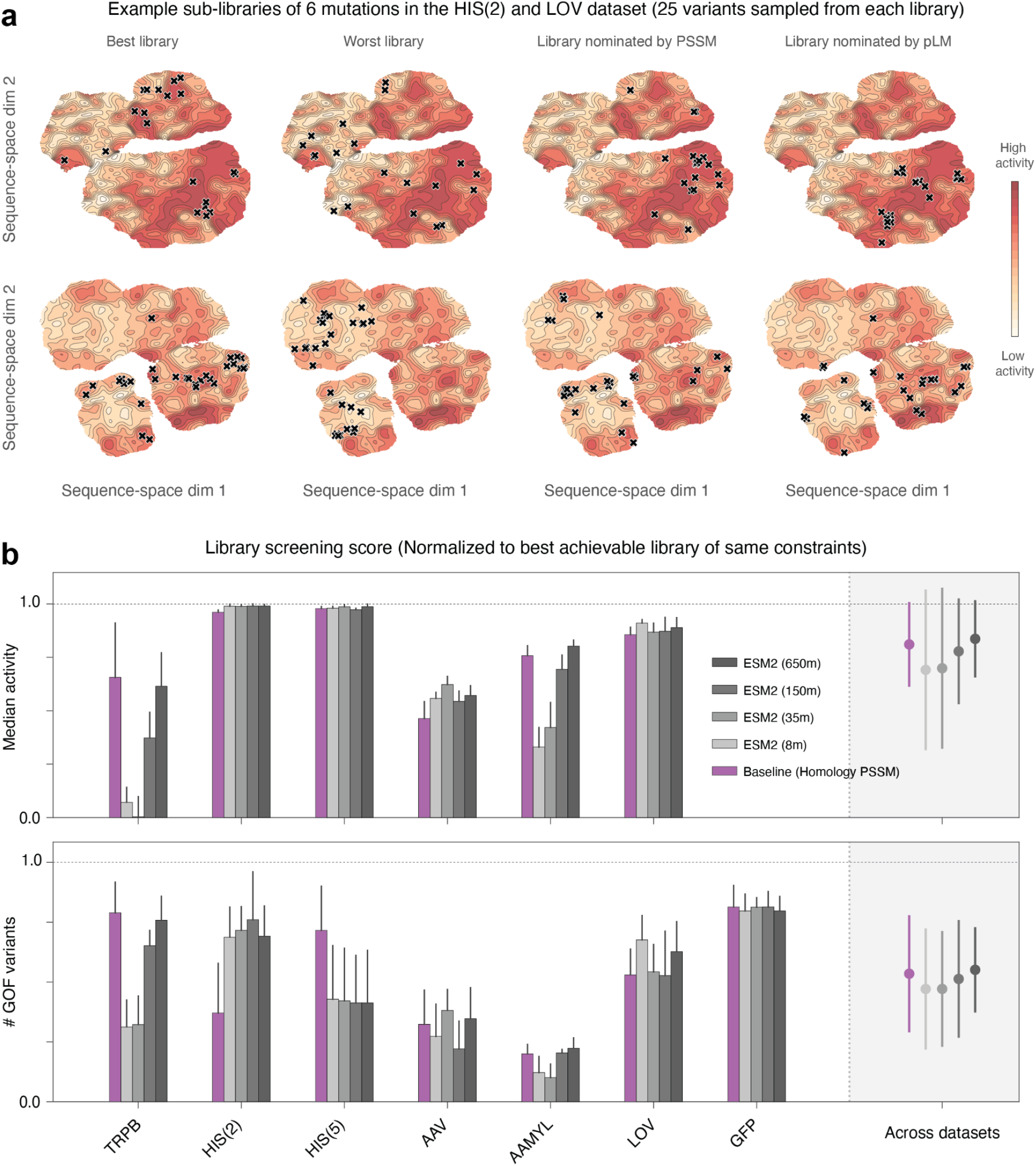
PSSM rivals pLMs in sequence-landscape functional selections. **(a)** Representative examples of model-guided sub-library nomination in two empirical fitness landscapes. Examples of 6-mutation sub-libraries are shown for the HIS(2) and LOV datasets, with 25 experimentally assayed variants sampled from each nominated library. For each dataset, the best-empirical and worst-empirical reference under the same constraints are shown alongside libraries nominated by a homology-based PSSM and by a pLM. Background contours indicate empirical activity across the plotted sequence space, and crosses denote experimentally assayed variants belonging to the corresponding nominated library. These landscape visualizations are shown for illustration only; all library selections were made directly from the underlying sequence-function data rather than from the plotted two-dimensional representation.**(b)** Summary of model-guided library-screening performance across empirical landscapes. For each dataset, each model nominated a sub-library defined by selected mutations across the designed positions, and experimentally measured variants belonging to that library were evaluated. Performance is summarized using two metrics: median activity of variants recovered in the nominated library (top) and the number of gain-of-function (GOF) variants recovered (bottom). Scores are normalized to the best empirical reference identified under the same design constraints within each empirical landscape, allowing comparison across datasets. Bars show model performance for individual datasets, and points at right summarize performance across datasets. Across the examined landscapes, PSSM-based libraries were generally comparable to those nominated by pLMs, and increasing pLM scale did not confer a clear or consistent advantage.

We quantified library quality using two complementary measures: the median activity of sampled variants and the number of gain-of-function variants recovered (Supplementary Fig. 18b). Because the landscapes differ in their activity levels and the prevalence of gain-of-function variants, we normalized each measure to the best empirical reference identified under the same design constraints. Across the densely sampled combinatorial landscapes that form the core of our benchmark, PSSM-nominated libraries were comparable to those nominated by pLMs, and larger pLMs did not yield consistent improvements. In some datasets, smaller pLMs performed as well as, or better than, larger models.

A modest apparent improvement with larger pLMs was observed when results were considered across all datasets, but this trend was driven largely by TrpB and AlphaAmyl, which differ from the densely sampled combinatorial landscapes emphasized in our benchmark. Within the mutation-dense landscapes, there was no consistent relationship between model size and library quality. Thus, although larger pLMs can be beneficial in specific cases, we did not observe a general scaling advantage for combinatorial library design.

In the main analysis, eligible variants could contain up to one mutation outside the set nominated by each method. To test whether this allowance influenced library quality, we repeated the analysis using only variants whose mutations were entirely contained within the nominated set (Supplementary Fig. 19b). The results remained qualitatively unchanged, with PSSM-nominated libraries performing comparably to those nominated by pLMs.

Taken together, these analyses indicate that pLMs can identify regions of sequence space enriched for functional variants, but they do not consistently outperform a simple evolutionary baseline. This result extends our zero-shot observations from individual variant scoring to a model-guided library-design setting and again shows that large pLMs do not reliably improve performance in the mutation-dense combinatorial landscapes.

## Discussion

Several studies have suggested that pLMs may serve as general-purpose tools for protein design^4,44^, motivated by their versatility and success in zero-shot scoring and transfer learning. But few publicly available datasets densely sample the combinatorial sequence spaces commonly explored in protein engineering, where candidate substitutions are combined to identify improved variants. We focused on this setting using controlled extrapolation from lower to higher mutational orders. Across these landscapes, predictive performance depended strongly on the mutational regimes represented in training and testing, while differences among pLM architectures and sizes were inconsistent. Surprisingly, pLMs did not consistently outperform simple mutation-identity or evolutionary baselines, including in model-guided combinatorial library selection. These findings do not imply that pLMs are uninformative for other protein-modeling tasks, but suggest that their advantages are limited in the design-of-function settings examined here.

Our findings also suggest that pretraining does not necessarily provide a superior representation for these tasks. One-hot encoding (OHE) remained competitive with, and often outperformed, pLM representations under both controlled extrapolation and random train-test splits. Within these combinatorial landscapes, much of the variation in pLM embeddings was recoverable from explicit mutation identity, while combining pretrained representations with mutation-identity or evolutionary features provided only small and inconsistent predictive gains. This is notable given the disparity in model complexity and computational requirements: the largest sequence-based pLM contains 3 billion parameters, and required computationally intensive pretraining^53,54^, whereas OHE requires no pretraining and simply records substitutions at experimentally varied positions. The benefit of pretrained representations may nevertheless depend on the prediction task. Li et al.^79^ reported advantages for pLM representations across several tasks, including those involving sequences from multiple protein families, where sequence variation extends far beyond a restricted set of substitutions. By contrast, the design problems examined here concern combinations of mutations within individual proteins, where much of the sequence remains fixed. This distinction may help explain why pretrained representations contribute limited additional predictive information in our benchmark, without implying that they are not useful for other protein-modeling tasks.

These observations also suggest a possible explanation for the strong influence of training-data composition. Our zero-shot analyses found that the evaluated pLM scoring procedures did not reliably reflect experimentally measured functional epistasis, raising the possibility that functionally relevant interactions must be learned primarily from experimental measurements. This could explain why including higher-order variants in training improves extrapolation even with simple mutation-based representations. Under this hypothesis, predictive performance may depend not only on training-set size or mutational order, but also on how well the measured variants capture the functional interactions relevant to the test sequences. Although consistent with our observations, this explanation remains to be tested directly.

Our conclusions concern primarily sequence-based pretrained pLMs, together with a structure-aware SaProt analysis and a focused evaluation of LoRA adaptation. They do not necessarily extend to homolog-conditioned or other multimodal approaches, which may incorporate additional target-specific information at inference. Given our observations, we stress that such models should be compared with baselines that use comparable information. Because homolog-conditioned pLMs use target-family sequences at inference, their performance should be compared with PSSMs or other baselines built from the same homolog information. Practical considerations also differ: PSSMs require a homolog search and alignment for each protein family, whereas pretrained pLMs avoid this requirement but incur high model inference costs. Their relative usefulness may therefore depend on homolog availability, the number of protein families being evaluated, and computational resources. These considerations warrant direct evaluation rather than assumptions based on model complexity.

More broadly, our results emphasize the importance of evaluating protein-design methods against simple, domain-informed baselines under experimentally relevant conditions. The central challenge in settings that are relevant to design of improved function is to identify favorable combinations of mutations whose functional consequences may be strongly context dependent. Increasing pLM scale or representation complexity alone did not consistently improve this task. Whether more effective integration of evolutionary, structural, or biophysical information can provide additional benefits remains an important question for future work^29,48,80,81^. Such improvements should ultimately be demonstrated against strong mutation-identity and evolutionary baselines, both in predicting unmeasured variants and in prioritizing experimentally useful combinatorial libraries.

## Online Methods

### Datasets and benchmark design

The principal benchmark comprised three FuncLib/htFuncLib landscapes, GFP, PTE and NMT^61^, and four ProteinGym assays, LOV, PARD3, GCN4 and HIS (Supplementary Fig. 1). The two combinatorial regions of HIS were analyzed separately as HIS(2) and HIS(5). AAV from FLIP was included where its sampling supported the same analysis, and TrpB and AlphaAmyl were additionally included in mutation-ranking and model-guided screening analyses. Models were compared on matched train/test partitions within each dataset. For GCN4 and PARD3, substitutions observed in fewer than seven variants were removed, followed by variants containing none of the retained substitutions.

### Protein language models

We benchmarked eight pretrained pLMs spanning multiple architectures, objectives and parameter scales: ESM2-8M, ESM2-35M, ESM2-150M, ESM2-650M, ESM2-3B, ProtBert-420M, ProGen2-small-170M and ProGen2-medium-700M (Supplementary Table 2). Public checkpoints were used throughout. ProtBert was excluded from zero-shot analyses because a compatible pretrained language-model output head was unavailable in our implementation.

We additionally evaluated SaProt-650M^58^ using the westlake-repl/SaProt_650M_AF2 checkpoint. For each dataset, 3Di tokens were generated from a reference structure using Foldseek structureto3didescriptor. The structure-derived sequence was aligned to the experimental wild type, positions lacking structural coordinates were assigned the SaProt missing-structure token, and, when several structural records were available, the record maximizing sequence identity times structural coverage was retained. The resulting 3Di sequence was fixed across all variants, with each variant amino-acid sequence interleaved with the same reference-derived structural tokens. Thus, SaProt tested the contribution of reference structural information rather than mutation-induced structural changes. Reference structures are listed in Supplementary Table 1.

### Landscape sampling geometry and phenotype definitions

Mutational order was defined as the number of amino-acid substitutions from the corresponding wild-type sequence. Variants were grouped by exact mutational order for Supplementary Fig. 1a,b. To quantify mutation reuse, we calculated the fraction of measured variants containing each observed substitution and compared these distributions with representative FLIP2 datasets. Pairwise sequence distance was calculated as Hamming distance over experimentally varied positions. For datasets containing more than 10,000 variants, distance distributions were estimated from 30 random samples of 5,000 variants rather than by enumerating all sequence pairs. Multidimensional scaling of Hamming distances was used only for sequence-space visualization.

Quantitative phenotypes were analyzed without thresholding. For classification analyses of continuous assays, variants with activity above the dataset median were labeled positive and variants at or below the median negative. GFP retained its original experimental binary labels, with activity defined as one minus the reported inactive label. The phenotype measured in each dataset and the corresponding positive fractions are summarized in Supplementary Table 1 and Supplementary Fig. 1b. Gain of function in the library-screening analyses was defined as activity above the dataset-specific 90th percentile.

### Zero-shot scoring

Zero-shot scores were calculated directly from the pretrained language-model output head without fitting experimental phenotypes. For masked language models, we used fixed-context additive and contextual, background-dependent scoring. For the fixed-context score, all experimentally variable positions *V* were masked simultaneously using the same input for every variant in a landscape. For substitution *i*,

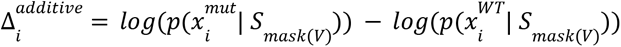

Because the input was shared across variants, each substitution received a fixed contribution. For a variant *S* containing mutation set *M*(*S*),

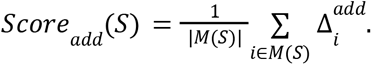

For contextual scoring, only positions mutated in the evaluated variant were jointly masked.

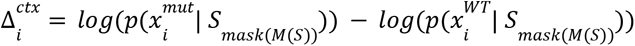

Experimentally variable positions that remained wild type were therefore visible, causing the conditioning context to vary between variants. The sequence score was again obtained by combining the mutant-versus-wild-type log-likelihood differences over the mutated positions. Because all mutated positions were masked simultaneously, however, the identities of jointly masked mutant residues did not directly condition one another. As a third scoring procedure, we evaluated each mutated position separately rather than masking multiple positions simultaneously. Each individual mutation was scored using its mutant-versus-wild-type log-likelihood difference in an independent forward pass, and the resulting mutation-level contributions were summed across the variant. This provided an additional additive zero-shot control in which each substitution was evaluated individually (Supplementary Fig. 4).

For ProGen2, scores were computed autoregressively rather than through masked-token prediction. Mutation-specific log-likelihood differences were calculated using the preceding sequence as context. Under fixed-context scoring, the preceding residues were taken from the wild-type sequence, whereas under background-dependent scoring they were taken from the evaluated variant, allowing upstream mutations to influence scores at downstream positions. Mutation-level contributions were then combined into a sequence-level score.

### Homology-based evolutionary baseline

As an evolutionary baseline, position-specific scoring matrices (PSSMs) were generated as described previously^36,37^ Sequences were clustered using CD-HIT^82^, aligned using MUSCLE^83^, and the resulting multiple sequence alignment was converted to a PSSM using PSI-BLAST^84^.

### Experimental epistasis

We quantified experimental pairwise epistasis for mutation pairs with measured wild-type, single-mutant, and double-mutant phenotypes. For mutations A and B, epistasis was defined as

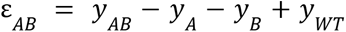

where *y* denotes the experimentally measured phenotype. We restricted pairwise analyses to experimentally measured double mutants for which both constituent single mutants were also available. To assess whether pLM scoring captured quantitative functional epistasis, we calculated the signed difference between contextual zero-shot scores and additive scores obtained by evaluating each mutation independently in the wild-type background, and correlated this difference with experimental pairwise epistasis using Spearman ρ separately for each model and dataset. This analysis assesses whether pLM scoring-context dependence reflects functional epistasis, rather than directly measuring mutation-identity-conditioned interactions.

For the complementary analysis of qualitative epistasis, continuous phenotypes were binarized using the dataset median, while GFP retained its original experimental labels. Expected double-mutant phenotype was defined using an AND rule: positive-positive pairs were expected to be positive, whereas positive-negative and negative-negative pairs were expected to be negative. Positive-positive pairs observed as negative were classified as negative sign epistasis, negative-negative pairs observed as positive as strong positive sign epistasis, and positive-negative pairs observed as positive as weak positive sign epistasis. Pairs matching the expected phenotype were retained as non-epistatic controls. This phenotype-class definition differs from conventional quantitative measures of epistasis, which define interactions as deviations from an assumed additive or multiplicative expectation. We used this coarse-grained formulation because it requires fewer assumptions about the scale and calibration of the underlying activity measurements and instead identifies pairs whose qualitative behavior is inconsistent with that expected from their constituent single mutants.

Because the expectation for positive-negative pairs is less constrained, analyses were also repeated after excluding these pairs (Supplementary Fig. 5,8-9). Higher-order variants were then examined for cases in which the same epistatic double-mutant combination was retained, but additional background mutations changed the binary phenotype relative to the measured double mutant. We classified such cases as higher-order reversals.

### Supervised representations

For each sequence, we extracted residue-level hidden representations from the final pLM layer and restricted them to the experimentally varied positions. Let *H^P^*∈ ℝ^mxd^ denote the representation at the *m* designed positions. Before downstream fitting, H*^P^* was L2-normalized across the position axis independently for each embedding dimension. Position-specific vectors were either averaged,

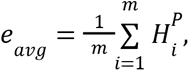

or flattened, *e_flat_* = *vec*(*H^p^*). Mean-pooled representations were subsequently Z-scored. Mean and flattened encodings were both considered during model selection, with mean pooling generally performing better. Alternative preprocessing, including normalization across the embedding dimension, pooled-vector L2 normalization and unnormalized mean pooling, gave similar results in representative controlled-extrapolation analyses. The one-hot encoding (OHE) baseline explicitly represented the amino-acid identity at each experimentally varied position and contained no pretrained information. OHE and pLM representations underwent independent downstream model selection so that each representation could use an appropriate predictor for its dimensionality.

### Evaluation protocols and primary downstream model selection

Controlled-extrapolation models were trained through a maximum mutational order *k_train_* and evaluated at higher exact test orders *k*_test_> *k_train_*. Results were analyzed either by fixing the test order and varying the training regime or by fixing the training regime and increasing the test order. Random train/test analyses were repeated for 50 iterations.

The primary benchmark used scikit-learn MLPRegressor and MLPClassifier. Model selection was performed independently for each dataset and representation using single- and double-mutant data, after which one architecture was selected for each dataset-representation combination.

The search considered single hidden layers of 20, 50, 100 or 200 units and two-layer architectures of 100-20 or 200-20 units; Adam and L-BFGS optimization; regularization strengths of 0.5, 0.1 and 0.01; Adam learning rates of 10^-4^and 10^-4^; and 25, 50, 100 or 500 maximum iterations. Mean and flattened pLM representations were included in this selection.

For controlled extrapolation, the selected architecture was held fixed across all lower-to-higher mutational-order comparisons. The same architecture was subsequently used for the corresponding random-split experiments and held fixed across random training-set sizes. Linear, ridge and gradient-boosted predictors were also examined during model development, with MLPs performing best overall.

### Statistical analyses and evaluation metrics

Model performance was evaluated using Spearman correlation for quantitative phenotypes and ROC AUC for classification. For zero-shot classification, ROC AUC was calculated in an orientation-invariant manner to measure separation between phenotype classes irrespective of score direction, whereas supervised classifiers used standard ROC AUC based on the predicted positive-class score. Precision@100 was defined as the fraction of positive variants among the 100 highest-ranked predictions, or among all test variants when fewer than 100 were available. Representation reconstruction was evaluated using R^2^

To quantify the association between predictive performance and benchmark factors, we calculated the factor-specific sum of squares relative to the total sum of squares. Factors included pLM identity, parameter-count bin (<50M, 50-500M, 500-700M, >700M), training mutational order and evaluation metric; training-set size was additionally included for GCN4 and NMT. These quantities were calculated separately for each factor and were not interpreted as mutually exclusive variance components.

Statistical significance was assessed using an F statistic with ANOVA where applicable. Otherwise, group labels were permuted 5,000 times and the empirical *P* value was calculated as the fraction of permuted F statistics exceeding the observed value.

### Alternative model selection procedure

We independently repeated the OHE-pLM comparison using a larger PyTorch architecture search with one to three hidden layers of 16 to 1,024 units, learning rates of 3 × 10^-4^to 10^-3^, weight decay of 10^-5^or 10^-4^, and dropout from 0 to 0.2. Candidate grids were adjusted to training-set size.

We tested three controlled-extrapolation selection strategies: (1) An internal validation within the available lower-order training data training through order *K*, (2) validation on order *K* + 1, and testing only above *K* + 1, and (3) an explicitly post-hoc procedure in which architecture selection used resampling across the broader dataset before refitting on the appropriate lower-order data. The latter was used as an optimistic oracle-like analysis rather than a prospective procedure. Random-split analyses similarly compared architecture selection within each sampled training set with broader post-hoc selection.

Architectures were selected by ROC AUC for classification and Spearman correlation for regression, except when a next-order regression validation set contained a single sequence, in which case mean squared error was used. A coarse search was followed by a finer search over the three best candidates. When validation scores differed by less than 0.01, the simpler architecture and earlier checkpoint were preferred. These analyses were used only as sensitivity tests (Supplementary Fig. 14).

### LoRA fine-tuning

We additionally fine-tuned ESM2-150M using LoRA implemented in PyTorch. LoRA rank was 16, alpha 32 and dropout 0.05, and a 64-unit prediction head was trained jointly. Optimization used AdamW with learning rate 3 × 10^-5^, weight decay 0.01, cosine decay, 10% warm-up, gradient clipping at 1.0, batch size 16, gradient accumulation of two, and up to 30 epochs. Evaluation batch size was 64 and seed 0 was used.

For a training cutoff *K*, models were fit on variants through order *K*. Twenty percent of order *K* + 1 was used for extrapolative validation, with the remaining *K* + 1 variants and all higher orders held out. Checkpoints were selected using Spearman correlation for regression or ROC AUC for classification, with patience of five epochs.

### Pooling and delta-embedding sensitivity

For NMT and PARD3, we additionally compared representations across transformer layers and sequence positions. Layer representations were combined by averaging, selecting individual layers or learned attention; positions were combined by global mean pooling, flattening or learned attention. Delta embeddings were calculated by subtracting the mean wild-type hidden-state tensor before pooling.

All pLM configurations used a 128-unit MLP with dropout 0.1 and weight decay 10^-4^, trained with AdamW for up to 70 epochs at learning rate 10^-3^, batch size 32 and seed 42. Twenty percent of the available lower-order training pool was used for validation and all variants above *K* remained held out. OHE was evaluated independently using a broader MLP grid under the same validation scheme.

### Strictly additive mutation-identity baseline

To evaluate phenotype prediction using a strictly additive baseline, we fit ordinary linear regression directly to one-hot mutation-identity features without pairwise or higher-order terms. Continuous phenotype was used for quantitative assays and the 0/1 activity label for GFP; ROC AUC for GFP was calculated from the continuous predictions. Features were not standardized.

For controlled extrapolation, models were fit to all variants with mutational order ≤ *K* and evaluated both jointly above *K* and separately at each exact higher order. Random-split analyses used training sizes of 10, 20, 50, 100, 1,000 and 5,000 variants where available, with 10 repeats and seed 0. This analysis was restricted to variants with at most 10 mutations.

### Representation geometry and linear reconstruction

Representation geometry was compared using pairwise Hamming distances between OHE variants and cosine distances between pLM embeddings. In each of 30 repeats, 1,500 variants were sampled without replacement from a random-number generator initialized with seed 0, and Spearman correlation was calculated between the upper-triangular entries of the two distance matrices. Representative heatmaps displayed 100 variants after rank-normalizing distances for visualization.

Within each 1,500-variant sample, variants were divided 50:50 into training and held-out subsets using the same split for all representations. Ridge regression was fit in both directions, OHE to pLM and pLM to OHE, using a training-fitted StandardScaler followed by RidgeCV over α = 10 and 100 with two-fold internal cross-validation.

Reconstruction was evaluated on held-out variants using global and variance-weighted multi-output R^2^; global R^2^ was used for the main reconstruction summaries.

### Empirical mutation ranking

For each candidate mutation *m*, empirical mutation score was defined as

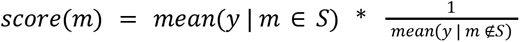

This score measures whether variants containing a mutation occupy, on average, a more favorable region of the experimentally measured landscape.

### Model-guided empirical sub-library screening

Each model ranked candidate substitutions, and the highest-ranked substitutions at distinct positions defined a nominated mutation set. In the main analysis, an experimentally measured variant was eligible if it contained at least one nominated substitution and at most one substitution outside the nominated set; nominated positions could remain wild type. Only experimentally measured eligible variants were sampled.

Main sub-library sizes were 30, 40 and 50 mutations for AlphaAmyl and 3-8 mutations for GFP, HIS(2), HIS(5), AAV, TrpB and LOV. Screening budgets were 5, 10, 25 and 50 variants. For each library-budget combination, eligible variants were sampled without replacement 30 times. No fixed seed was used for these resampling iterations. Library quality was measured by the mean number of gain-of-function variants recovered and the median across resamples of the within-sample median activity. Combinations with no more eligible variants than the requested budget were excluded.

Scores were normalized to an empirical reference identified under the same library-size, eligibility and screening constraints. Reference mutation sets were optimized separately for gain-of-function recovery and median activity using a fixed-size greedy swap search with six restarts, up to 12 rounds and up to 250 swaps per round. Candidate additions were drawn from the 40 highest-ranked, 40 highest-coverage and 40 random substitutions. Median-activity candidates used seven resamples during search, followed by reevaluation of the top 30 candidates with 500 resamples. This reference search used deterministic seeds derived from seed 0 and, because it was heuristic rather than exhaustive, is referred to as the best empirical reference identified by the search.

As a stricter sensitivity analysis, we required all substitutions in an eligible variant to belong to the nominated set. Strict sub-library sizes were 30, 40 and 50 for AlphaAmyl; 6-8 for GFP, HIS(2), HIS(5) and LOV; 8-10 for AAV; and 6-12 for TrpB. Screening budgets were 5, 10 and 25 for AlphaAmyl, GFP, HIS(2), HIS(5) and LOV; 5, 8 and 10 for AAV; and 5, 10 and 25 for TrpB. All other procedures were unchanged.

## Data Availability

All datasets used in this study, except the NMT dataset, are publicly available and the sources listed in Supplementary Table 2. The NMT dataset was provided by Lucas Bocquin and Stephan C. Hammer (Bielefeld University) and is described in a separate study.

## Code Availability

Code and notebooks are available at https://github.com/drprfitay/simple_baselines.

## Use of Generative AI

Generative AI tools were used as aids for scientific writing and programming. For writing, AI was used primarily to improve clarity, structure, concision, and phrasing of text whose scientific content, interpretation, and conclusions were provided by the authors. For programming, AI was used to assist with implementation of analyses specified and designed by the authors. All AI-assisted code was reviewed and validated by the authors, including through inspection, consistency checks, assertions, comparison with independently obtained or previously reproduced results, and/or repeated analyses where appropriate. AI was not used to generate scientific hypotheses, experimental results, data, or conclusions independently. The authors take full responsibility for all analyses, code, interpretations, and statements presented in the manuscript.

## Acknowledgements

We thank James Bowden (UC Berkeley), Dina Listov, Omri Porat, Tom Talpir, Ariel Tennenhouse, and Shlomo Yakir Hoch (Weizmann Institute of Science) for fruitful discussions and helpful comments on the manuscript. We thank Stephan C. Hammer and Lucas Bocquin (Bielefeld University) for generously providing access to the unpublished NMT dataset and for permission to analyze it in this study. The study was supported by a European Research Council Advanced Grant (101140394).

