## Supplemental figures for "Simple baselines rival protein language models in dense combinatorial protein landscapes"

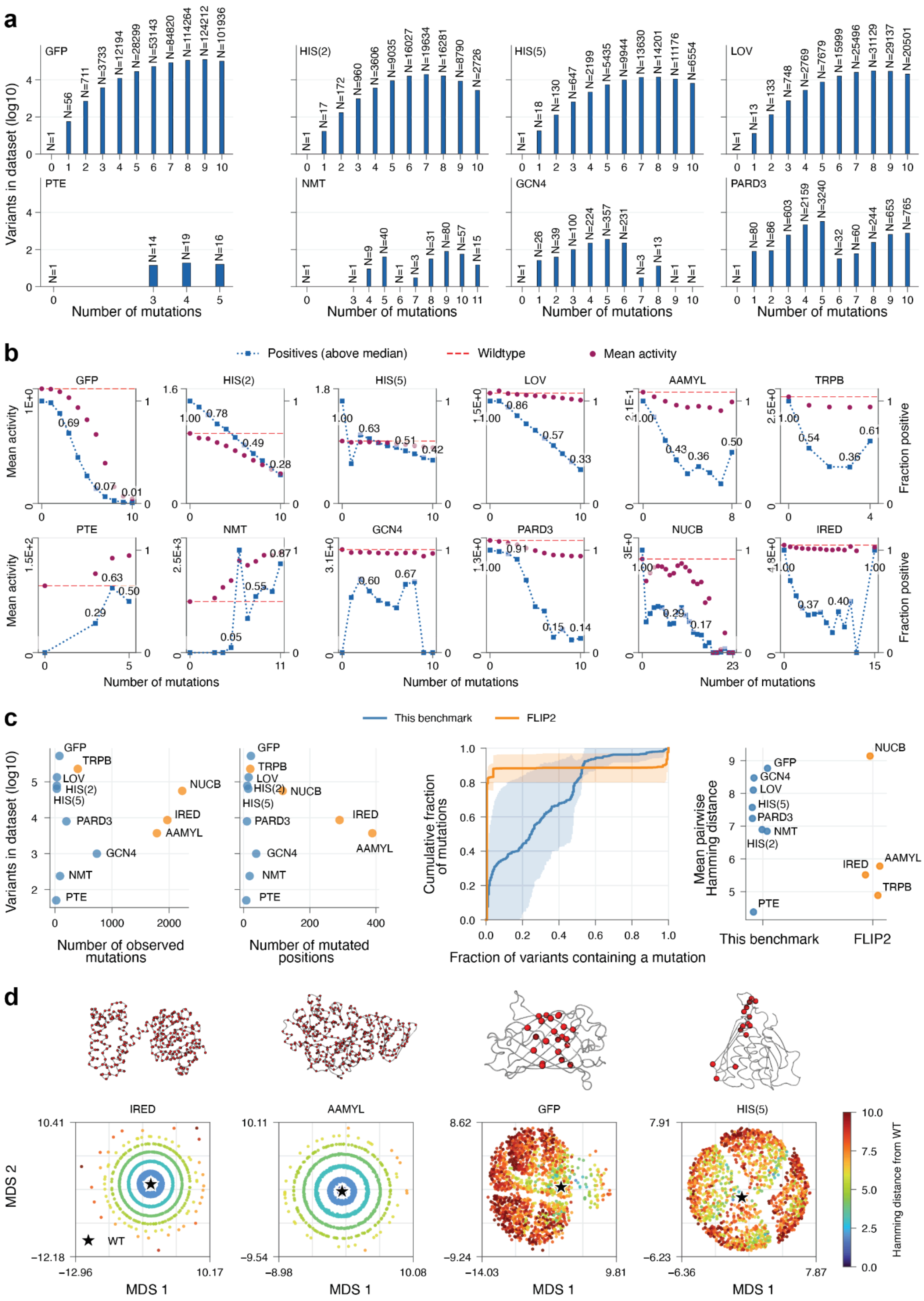

a
Variants in dataset (log10)
GFP
N=1
N=56
N=711
N=3733
N=12194
N=28299
N=53143
N=84820
N=114264
N=124212
N=101936
HIS(2)
N=1
N=17
N=172
N=960
N=3606
N=9035
N=16027
N=19634
N=16281
N=8790
N=2726
HIS(5)
N=1
N=18
N=130
N=647
N=2199
N=5435
N=9944
N=13630
N=14201
N=11176
N=6554
LOV
N=1
N=13
N=133
N=748
N=2769
N=7679
N=15999
N=25496
N=31129
N=29137
N=20501
PTE
N=1
N=14
N=19
N=16
NMT
N=1
N=1
N=9
N=40
N=1
N=3
N=31
N=80
N=57
N=15
GCN4
N=1
N=26
N=39
N=100
N=224
N=357
N=231
N=3
N=13
N=1
N=1
PARD3
N=1
N=80
N=86
N=603
N=2159
N=3240
N=32
N=60
N=244
N=653
N=765
Number of mutations
b
Positives (above median)
Wildtype
Mean activity
Mean activity
Fraction positive
GFP
HIS(2)
HIS(5)
LOV
AAMYL
TRPB
PTE
NMT
GCN4
PARD3
NUCB
IRED
Number of mutations
c
This benchmark
FLIP2
Variants in dataset (log10)
Number of observed mutations
Number of mutated positions
Cumulative fraction of mutations
Fraction of variants containing a mutation
Mean pairwise Hamming distance
This benchmark
FLIP2
GFP
TRPB
LOV
HIS(2)
HIS(5)
PARD3
GCN4
NMT
PTE
NUCB
IRED
AAMYL
d
IRED
AAMYL
GFP
HIS(5)
MDS 1
MDS 2
WT
Hamming distance from WT

**<u>Supplementary Figure 1 - Properties and sampling geometry of analyzed fitness landscapes</u>**

**(a)** Distribution of experimentally measured variants across mutational orders. Bars show the number of variants at each exact number of mutations from the corresponding wild-type sequence on a log10 scale; labels above bars reflect the underlying counts.

**(b)** Measured phenotype as a function of mutational order. Magenta points show the mean measured phenotype at each order. Where applicable, the secondary axis shows the fraction of positive variants, using the original binary GFP labels or an above-median label for continuous assays. (see Methods) The wild-type phenotype is shown for reference as the horizontal red-dashed line.

**(c)** Comparison of the sampling geometry of landscapes analyzed here with representative FLIP2 datasets (enzymes). Left, the number of measured variants as a function of the number of distinct observed substitutions and experimentally varied positions. Middle, cumulative distributions of substitution frequencies across measured variants. In dense combinatorial landscapes, mutations recur across a broad and relatively continuous range of variant frequencies, whereas in less combinatorial landscapes they tend to occur in either almost all or very few measured variants. Right, mean pairwise Hamming distance between measured variants. For datasets containing more than 10,000 variants, Hamming-distance statistics were estimated over 30 repeated samples rather than enumerating all sequence pairs.

**(d)** Experimentally varied residues are highlighted on reference structures. In the representative FLIP2 datasets, mutations are distributed broadly across the protein, whereas the landscapes analyzed here typically vary a restricted set of selected positions, often concentrated in functionally important or otherwise sensitive regions, and repeatedly sample combinations among those mutations. Sequence-space projections were generated from pairwise Hamming distances using multidimensional scaling (MDS); points are colored by mutational distance from the wild type, marked by a star.

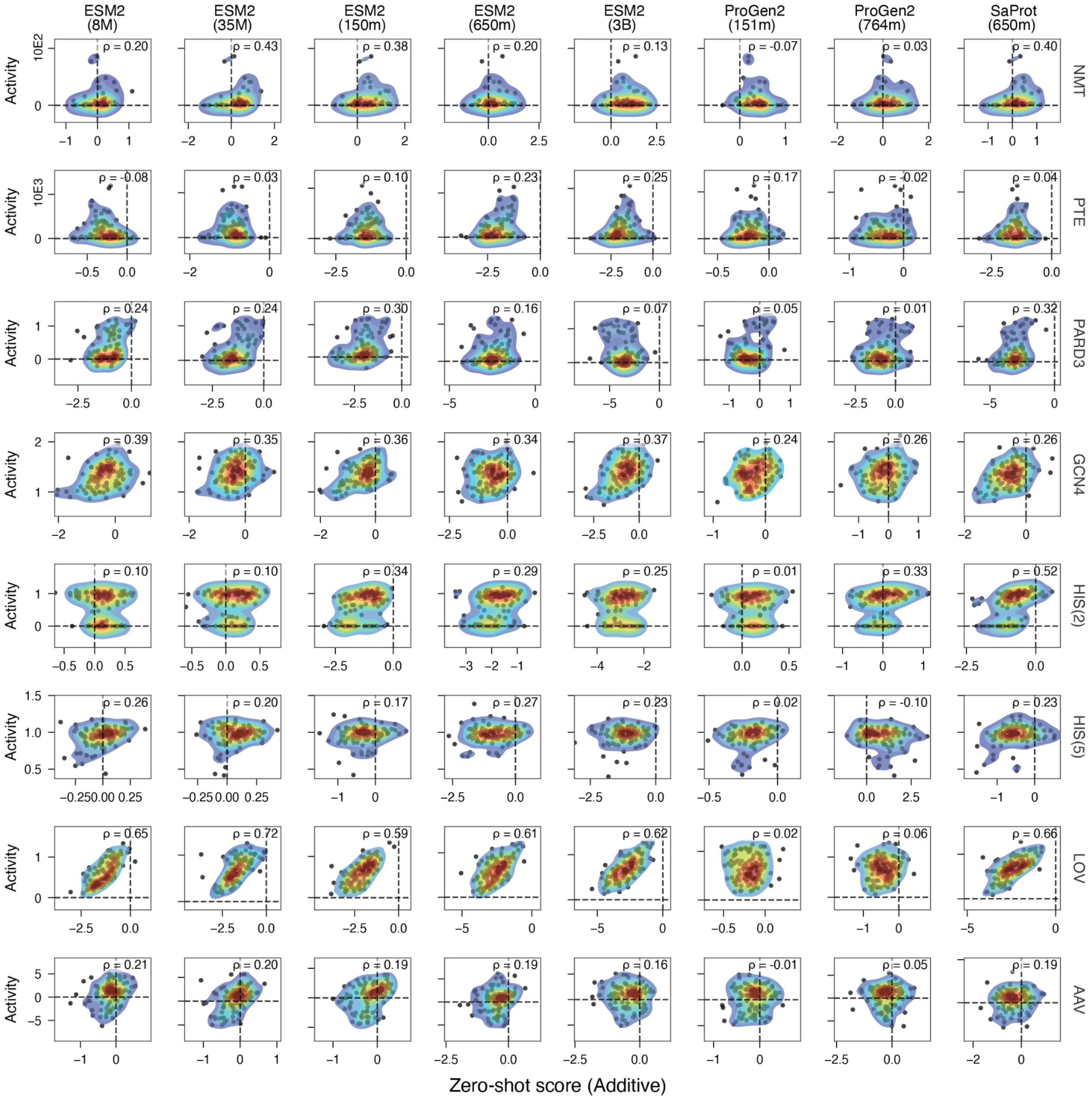
ESM2 (8M)
ESM2 (35M)
ESM2 (150m)
ESM2 (650m)
ESM2 (3B)
ProGen2 (151m)
ProGen2 (764m)
SaProt (650m)
Activity
10E2
10E3
NMT
PTE
PARD3
GCN4
HIS(2)
HIS(5)
LOV
AAV
ρ = 0.20 ρ = 0.43 ρ = 0.38 ρ = 0.20 ρ = 0.13 ρ = -0.07 ρ = 0.03 ρ = 0.40
ρ = -0.08 ρ = 0.03 ρ = 0.10 ρ = 0.23 ρ = 0.25 ρ = 0.17 ρ = -0.02 ρ = 0.04
ρ = 0.24 ρ = 0.24 ρ = 0.30 ρ = 0.16 ρ = 0.07 ρ = 0.05 ρ = 0.01 ρ = 0.32
ρ = 0.39 ρ = 0.35 ρ = 0.36 ρ = 0.34 ρ = 0.37 ρ = 0.24 ρ = 0.26 ρ = 0.26
ρ = 0.10 ρ = 0.10 ρ = 0.34 ρ = 0.29 ρ = 0.25 ρ = 0.01 ρ = 0.33 ρ = 0.52
ρ = 0.26 ρ = 0.20 ρ = 0.17 ρ = 0.27 ρ = 0.23 ρ = 0.02 ρ = -0.10 ρ = 0.23
ρ = 0.65 ρ = 0.72 ρ = 0.59 ρ = 0.61 ρ = 0.62 ρ = 0.02 ρ = 0.06 ρ = 0.66
ρ = 0.21 ρ = 0.20 ρ = 0.19 ρ = 0.19 ρ = 0.16 ρ = -0.01 ρ = 0.05 ρ = 0.19
Zero-shot score (Additive)

**Supplementary Figure 2 - Fixed-context additive zero-shot predictions across models and landscapes**

Fixed-context additive zero-shot predictions across the evaluated pLMs and compatible landscapes. Each panel compares experimental phenotype with the model score for one model-dataset combination, with Spearman's ρ shown in the panel. For masked language models, all experimentally variable positions were masked in a common sequence context for every variant, and mutant-versus-wild-type log-likelihood differences were combined only for substitutions present in that variant (see Methods). The conditioning input is therefore shared across variants and each substitution has a fixed contribution. Columns denote models and rows denote datasets.

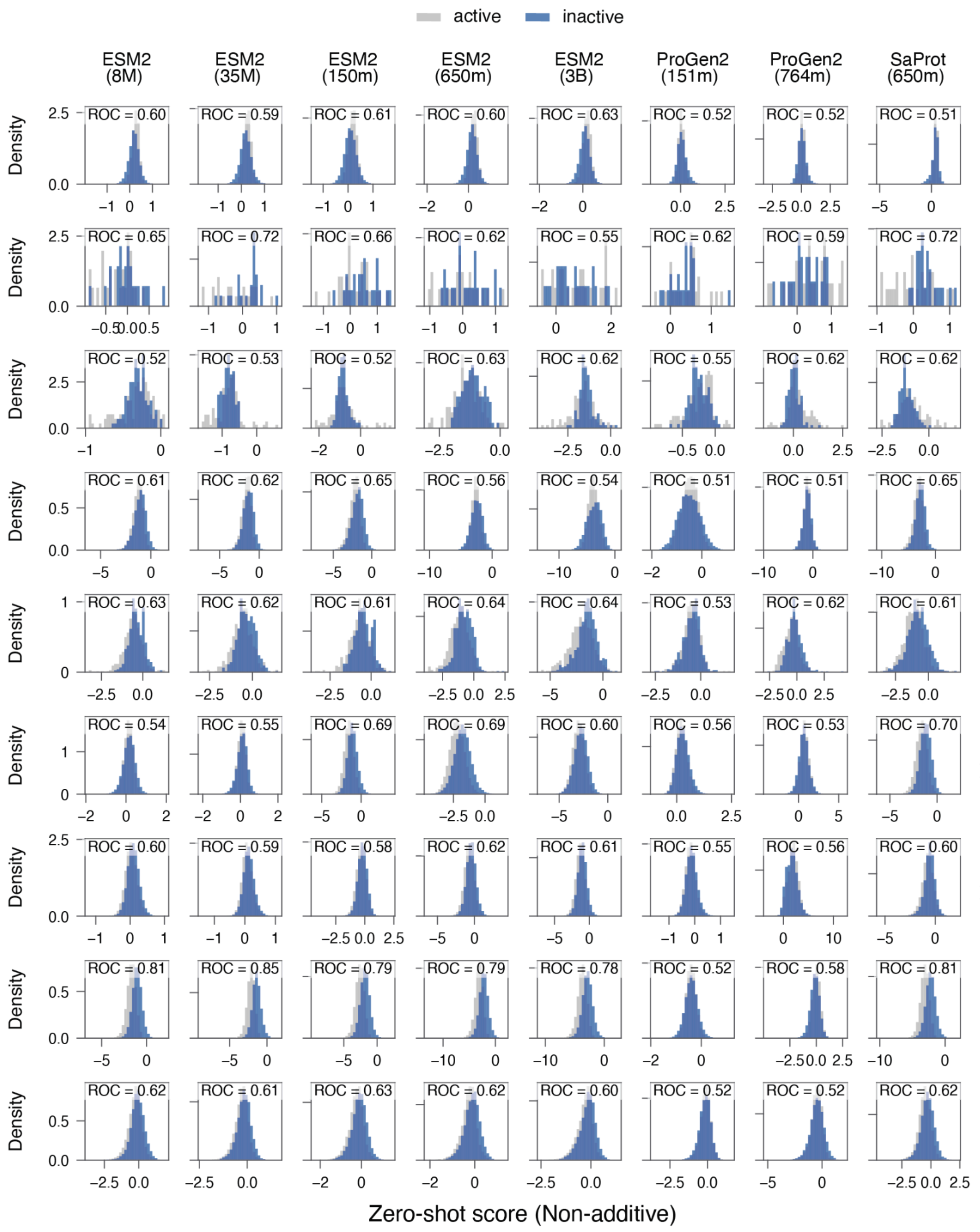
active
inactive
ESM2 (8M)
ESM2 (35M)
ESM2 (150m)
ESM2 (650m)
ESM2 (3B)
ProGen2 (151m)
ProGen2 (764m)
SaProt (650m)
GFP
ROC = 0.60
ROC = 0.59
ROC = 0.61
ROC = 0.60
ROC = 0.63
ROC = 0.52
ROC = 0.52
ROC = 0.51
NMT
ROC = 0.65
ROC = 0.72
ROC = 0.66
ROC = 0.62
ROC = 0.55
ROC = 0.62
ROC = 0.59
ROC = 0.72
PTE
ROC = 0.52
ROC = 0.53
ROC = 0.52
ROC = 0.63
ROC = 0.62
ROC = 0.55
ROC = 0.62
ROC = 0.62
PARD3
ROC = 0.61
ROC = 0.62
ROC = 0.65
ROC = 0.56
ROC = 0.54
ROC = 0.51
ROC = 0.51
ROC = 0.65
GCN4
ROC = 0.63
ROC = 0.62
ROC = 0.61
ROC = 0.64
ROC = 0.64
ROC = 0.53
ROC = 0.62
ROC = 0.61
HIS(2)
ROC = 0.54
ROC = 0.55
ROC = 0.69
ROC = 0.69
ROC = 0.60
ROC = 0.56
ROC = 0.53
ROC = 0.70
HIS(5)
ROC = 0.60
ROC = 0.59
ROC = 0.58
ROC = 0.62
ROC = 0.61
ROC = 0.55
ROC = 0.56
ROC = 0.60
LOV
ROC = 0.81
ROC = 0.85
ROC = 0.79
ROC = 0.79
ROC = 0.78
ROC = 0.52
ROC = 0.58
ROC = 0.81
AAV
ROC = 0.62
ROC = 0.61
ROC = 0.63
ROC = 0.62
ROC = 0.60
ROC = 0.52
ROC = 0.52
ROC = 0.62
Density
Zero-shot score (Non-additive)

**Supplementary Figure 3 - Contextual zero-shot classification across models and landscapes**

Zero-shot invariant ROC-AUC score of active and inactive variants using the contextual, background-dependent scoring procedure (see Methods). Histograms show score distributions for active and inactive variants for each model-dataset combination, and ROC AUC is reported in each panel. For masked language models, only positions mutated in the evaluated variant were masked, leaving experimentally variable positions that remained wild type visible to the model. ProGen2 contextual scores were obtained by left-to-right decoding, so that later positions were scored conditional on residues revealed at preceding positions. Columns correspond to models and rows to datasets.

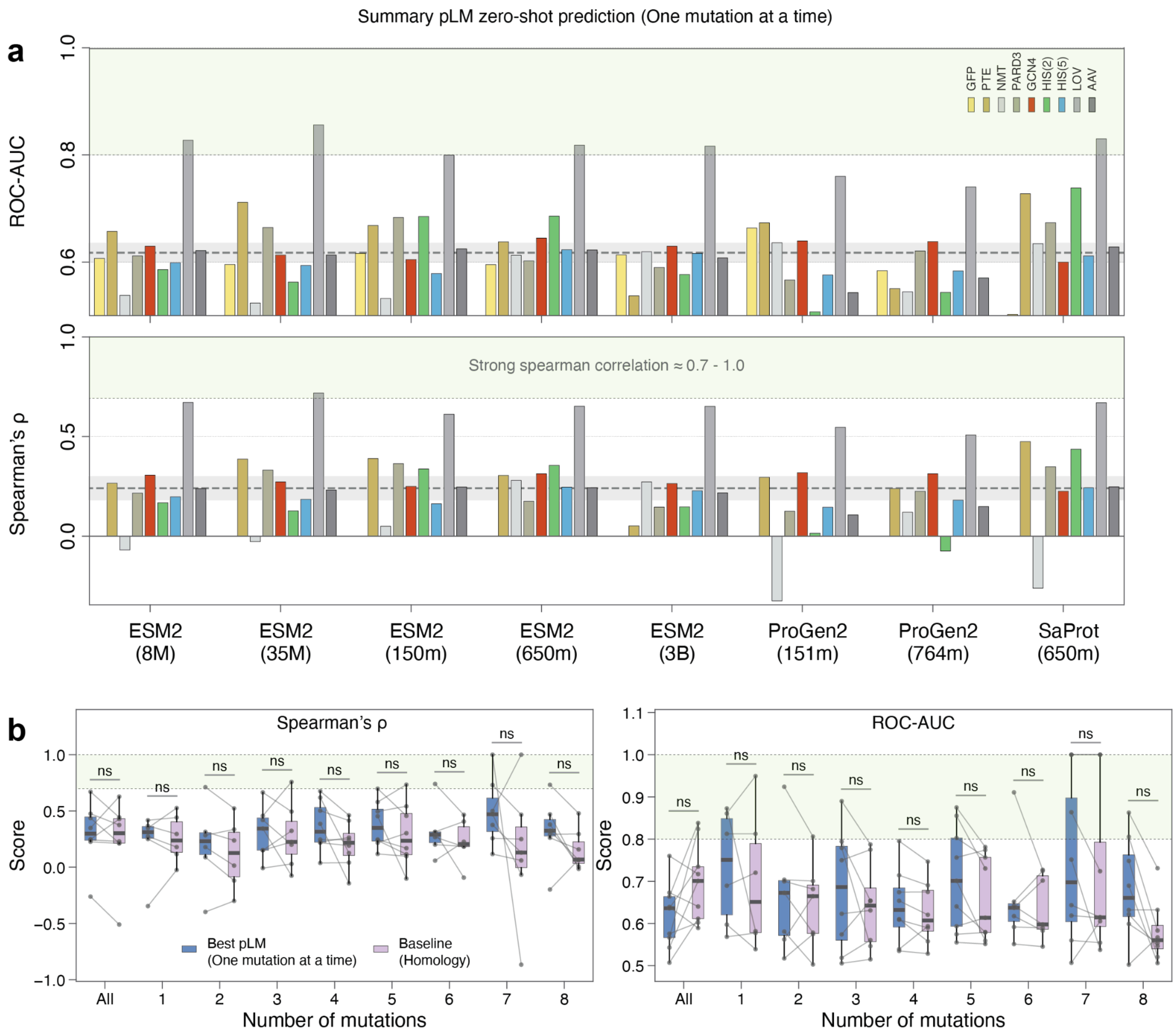
a
Summary pLM zero-shot prediction (One mutation at a time)
ROC-AUC
Spearman's ρ
GFP
PTE
NMT
PARD3
GCN4
HIS(2)
HIS(5)
LOV
AAV
Strong spearman correlation ≈ 0.7 - 1.0
ESM2 (8M)
ESM2 (35M)
ESM2 (150m)
ESM2 (650m)
ESM2 (3B)
ProGen2 (151m)
ProGen2 (764m)
SaProt (650m)
b
Spearman's ρ
ROC-AUC
Score
ns
Best pLM (One mutation at a time)
Baseline (Homology)
Number of mutations
All
1
2
3
4
5
6
7
8

**Supplementary Figure 4 - Independently masked additive zero-shot scoring**

**(a)** Zero-shot ROC AUC and Spearman correlation across compatible landscapes and pLMs when each substitution is scored separately in the wild-type background and mutation-level contributions are summed. Bars represent datasets.

**(b)** Comparisons between the best-performing pLM under this scoring procedure and the homology-based PSSM baseline, stratified by mutational order. Paired observations represent corresponding dataset comparisons; Differences are not significant (paired t-test; n.s., $P > 0.05$).

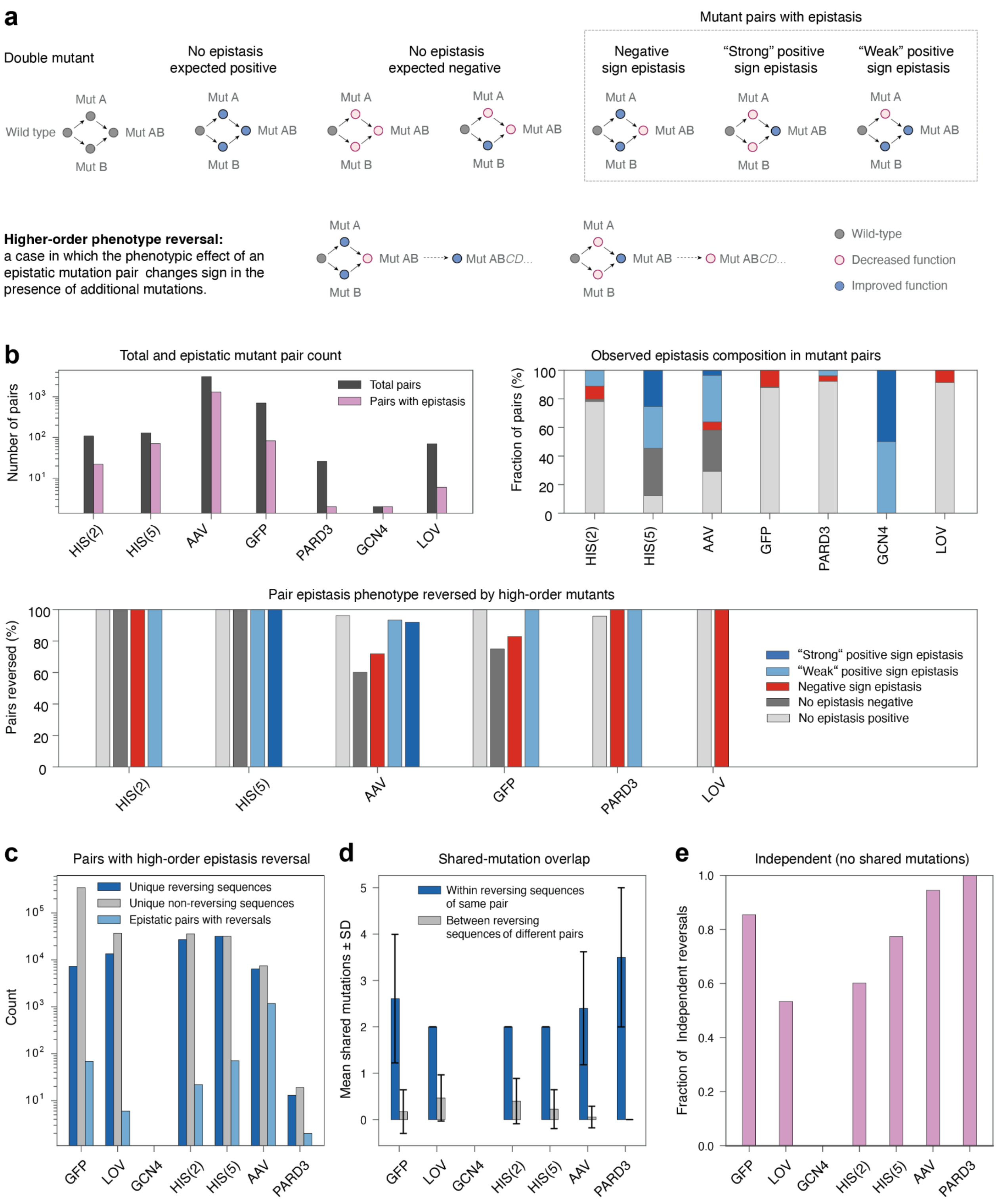

a
Mutant pairs with epistasis
Double mutant
No epistasis expected positive
No epistasis expected negative
Negative sign epistasis
"Strong" positive sign epistasis
"Weak" positive sign epistasis
Wild type
Mut A
Mut B
Mut AB
Higher-order phenotype reversal: a case in which the phenotypic effect of an epistatic mutation pair changes sign in the presence of additional mutations.
Mut ABCD...
Wild-type
Decreased function
Improved function
b
Total and epistatic mutant pair count
Number of pairs
Total pairs
Pairs with epistasis
Observed epistasis composition in mutant pairs
Fraction of pairs (%)
HIS(2)
HIS(5)
AAV
GFP
PARD3
GCN4
LOV
Pair epistasis phenotype reversed by high-order mutants
Pairs reversed (%)
"Strong" positive sign epistasis
"Weak" positive sign epistasis
Negative sign epistasis
No epistasis negative
No epistasis positive
c
Pairs with high-order epistasis reversal
Count
Unique reversing sequences
Unique non-reversing sequences
Epistatic pairs with reversals
d
Shared-mutation overlap
Mean shared mutations ± SD
Within reversing sequences of same pair
Between reversing sequences of different pairs
e
Independent (no shared mutations)
Fraction of Independent reversals

**Supplementary Figure 5 - Qualitative epistasis pairs and higher-order phenotype reversals**

**(a)** Definitions used to classify measured mutation pairs. Positive-positive single-mutant pairs were expected to be positive, whereas positive-negative and negative-negative pairs were expected to be negative. We classified a positive-positive pair that became negative as negative sign epistasis; a negative-negative pair that became positive as strong positive sign epistasis; and a mixed positive-negative pair that became positive as weak positive sign epistasis. We recorded a higher-order phenotype reversal when a measured variant containing an epistatic pair changed phenotype class after additional mutations were introduced.

**(b)** Top left, number of evaluable mutation pairs and number classified as epistatic in each landscape. Top right, composition of the observed epistasis classes. Bottom, fraction of pairs for which at least one higher-order phenotype reversal was observed, separated by epistasis class.

**(c)** Counts of unique higher-order variants that reversed or retained the measured pairwise phenotype, together with the number of epistatic pairs for which at least one higher-order reversal was observed.

**(d)** Mean overlap of additional background mutations among higher-order variants that reverse epistatic pairs. Within-pair overlap was calculated between reversing variants containing the same epistatic pair, whereas between-pair overlap was calculated between reversing variants associated with different epistatic pairs. Error bars show standard deviation. We used this comparison to characterize the complexity of higher-order epistatic reversals. High within-pair overlap indicates that reversal of a given epistatic pair tends to occur in similar mutational backgrounds, whereas low between-pair overlap indicates that different epistatic pairs are reversed by distinct sets of background mutations. Thus, for example, the pattern observed in LOV suggests that reversals are relatively stereotyped within individual epistatic pairs but diverse across different pairs.

(e) Fraction of higher-order reversals classified as independent, defined by the absence of a shared additional mutation with the other reversing backgrounds included in the comparison.

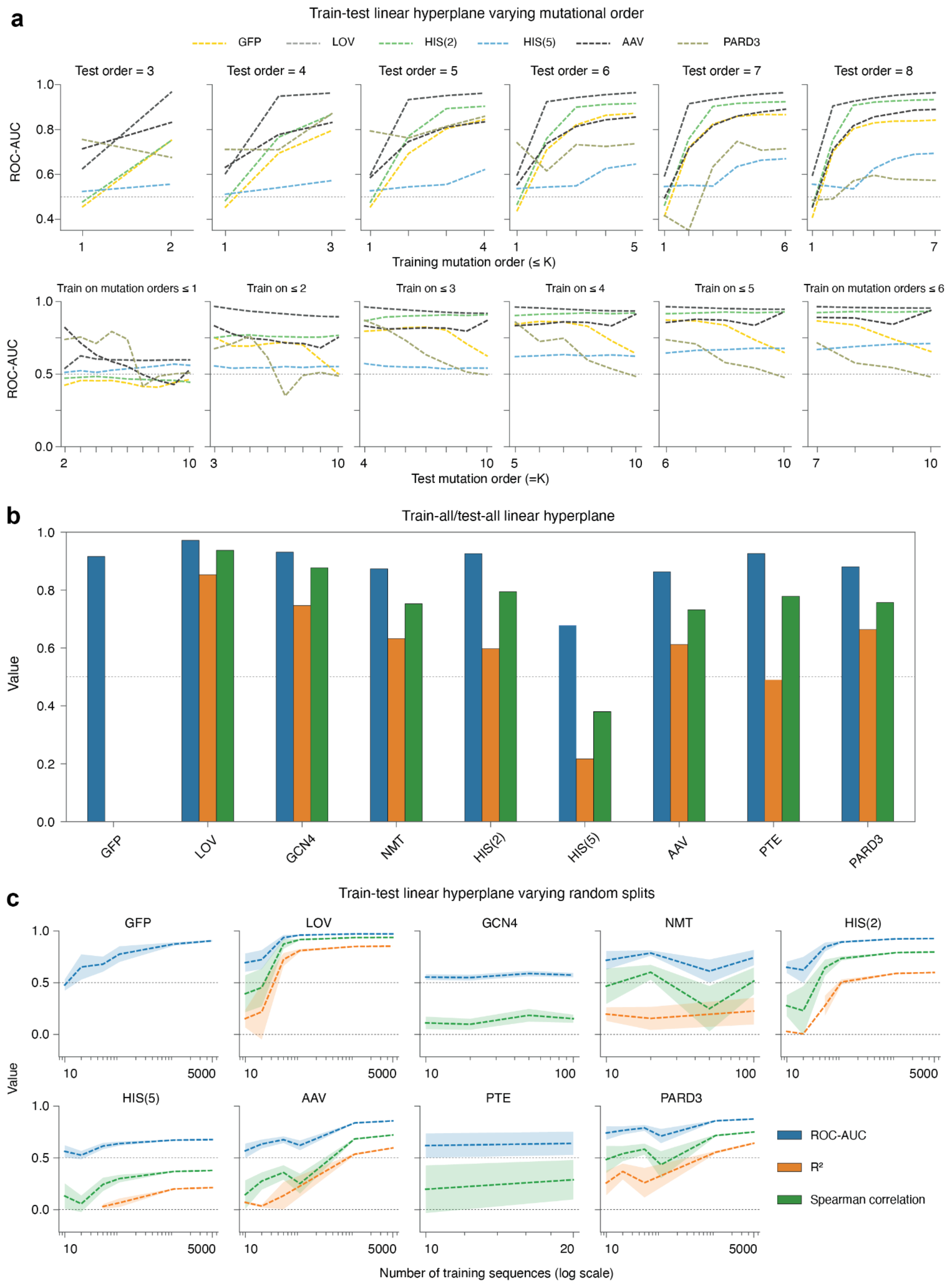
a
Train-test linear hyperplane varying mutational order
GFP
LOV
HIS(2)
HIS(5)
AAV
PARD3
Test order = 3
Test order = 4
Test order = 5
Test order = 6
Test order = 7
Test order = 8
ROC-AUC
Training mutation order (≤ K)
Train on mutation orders ≤ 1
Train on ≤ 2
Train on ≤ 3
Train on ≤ 4
Train on ≤ 5
Train on mutation orders ≤ 6
Test mutation order (=K)
b
Train-all/test-all linear hyperplane
Value
GFP
LOV
GCN4
NMT
HIS(2)
HIS(5)
AAV
PTE
PARD3
c
Train-test linear hyperplane varying random splits
GFP
LOV
GCN4
NMT
HIS(2)
HIS(5)
AAV
PTE
PARD3
Value
ROC-AUC
R²
Spearman correlation
Number of training sequences (log scale)

**Supplementary Figure 6 - Performance of additive mutation-identity predictors**

**(a)** Controlled mutational-order extrapolation using a strictly additive linear predictor fitted to mutation-identity features. Top, ROC AUC for fixed held-out test orders as progressively higher mutational orders are included in training. Bottom, the complementary analysis in which the training regime is fixed and performance is evaluated across increasingly higher-order test variants. Lines represent datasets, and the horizontal dotted line indicates chance classification (ROC AUC = 0.5). Performance is near chance in some datasets when training is restricted to single mutants, illustrating the difficulty of predicting combinations of individually tolerated mutations.

**(b)** Full-data diagnostic fits of the same additive predictor, trained and evaluated on the complete measured landscape. ROC AUC, $R^2$, and Spearman correlation are reported as applicable. These analyses assess how much of the observed phenotype variation can be captured by additive mutation-identity effects without explicitly modeling interactions between mutations.

**(c)** Performance of the additive predictor under conventional random train-test subsampling as a function of training-set size. Lines show mean performance across repeated random splits, and shaded regions indicate variation between splits. All predictors use mutation-identity features without interaction terms.

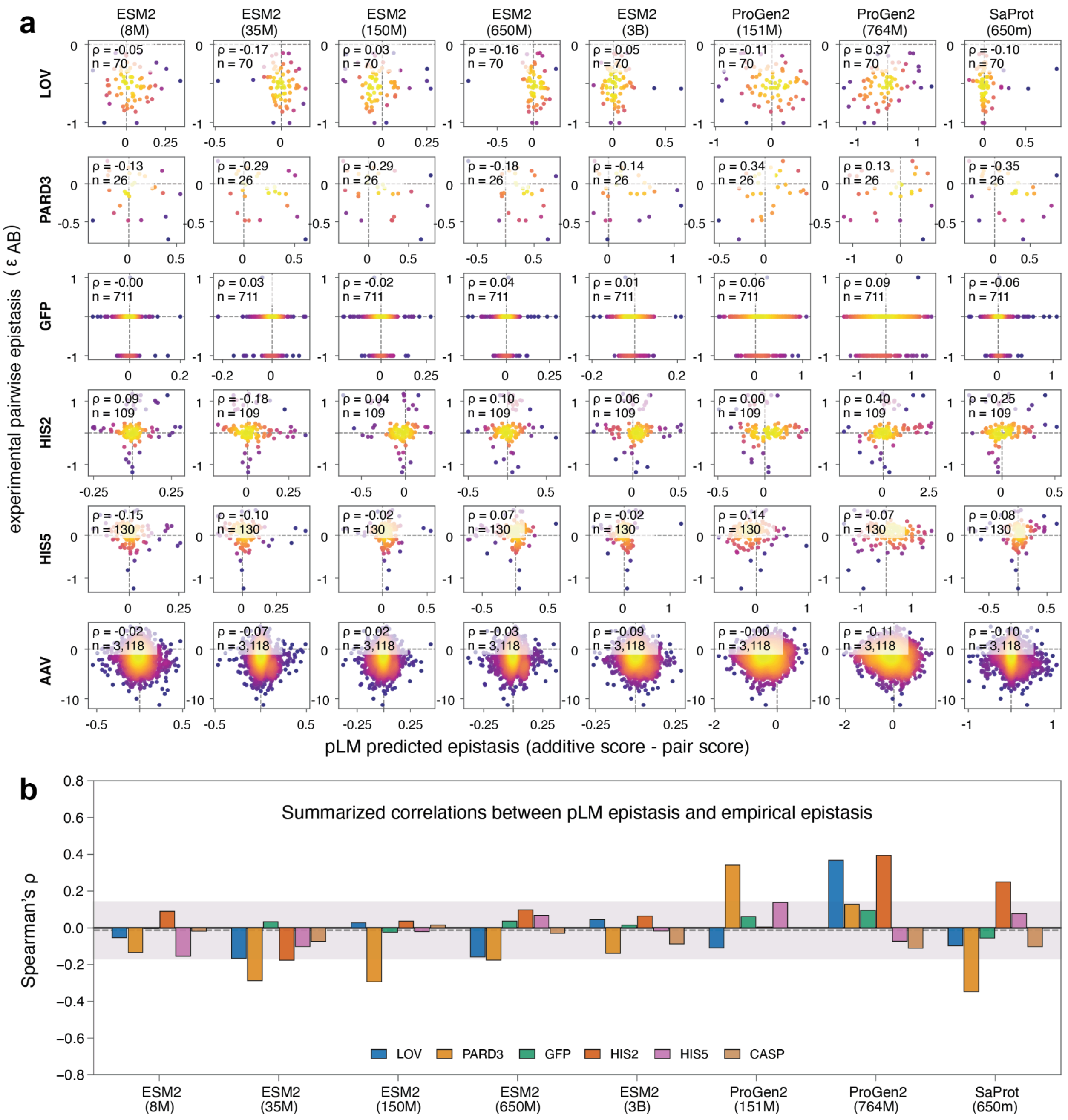
a
ESM2 (8M)
ESM2 (35M)
ESM2 (150M)
ESM2 (650M)
ESM2 (3B)
ProGen2 (151M)
ProGen2 (764M)
SaProt (650m)
LOV
PARD3
GFP
HIS2
HIS5
AAV
experimental pairwise epistasis (ε AB)
pLM predicted epistasis (additive score - pair score)
b
Summarized correlations between pLM epistasis and empirical epistasis
Spearman's ρ
LOV
PARD3
GFP
HIS2
HIS5
CASP

**Supplementary Figure 7 - Quantitative functional epistasis and pLM sensitivity to sequence background**

**(a)** Experimental pairwise functional epistasis (see Methods) plotted against the signed difference between contextual and independently masked additive zero-shot scores for measured double mutants with both constituent single-mutant phenotypes available. Rows show datasets and columns show pLMs. Each panel reports Spearman's ρ and the number of evaluated mutation pairs.

**(b)** Summary of Spearman correlations across model–dataset combinations. Correlations close to zero indicate little correspondence between the pLM score differences and experimentally measured functional epistasis.

a

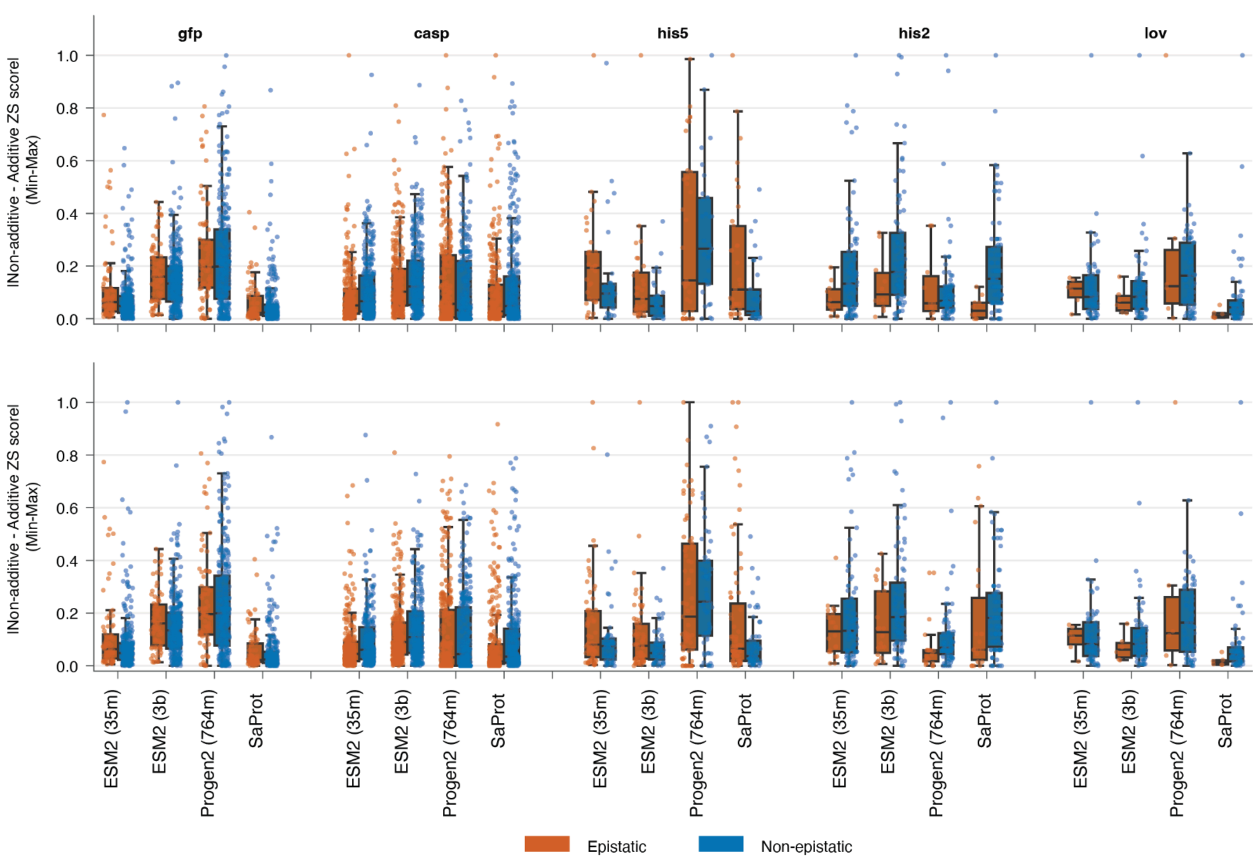
Strong positive + negative epistasis only
gfp
casp
his5
his2
lov
lNon-additive - Additive ZS scorel (Min-Max)
1.0
0.8
0.6
0.4
0.2
0.0
ESM2 (35m)
ESM2 (3b)
Progen2 (764m)
SaProt
Epistatic
Non-epistatic

b

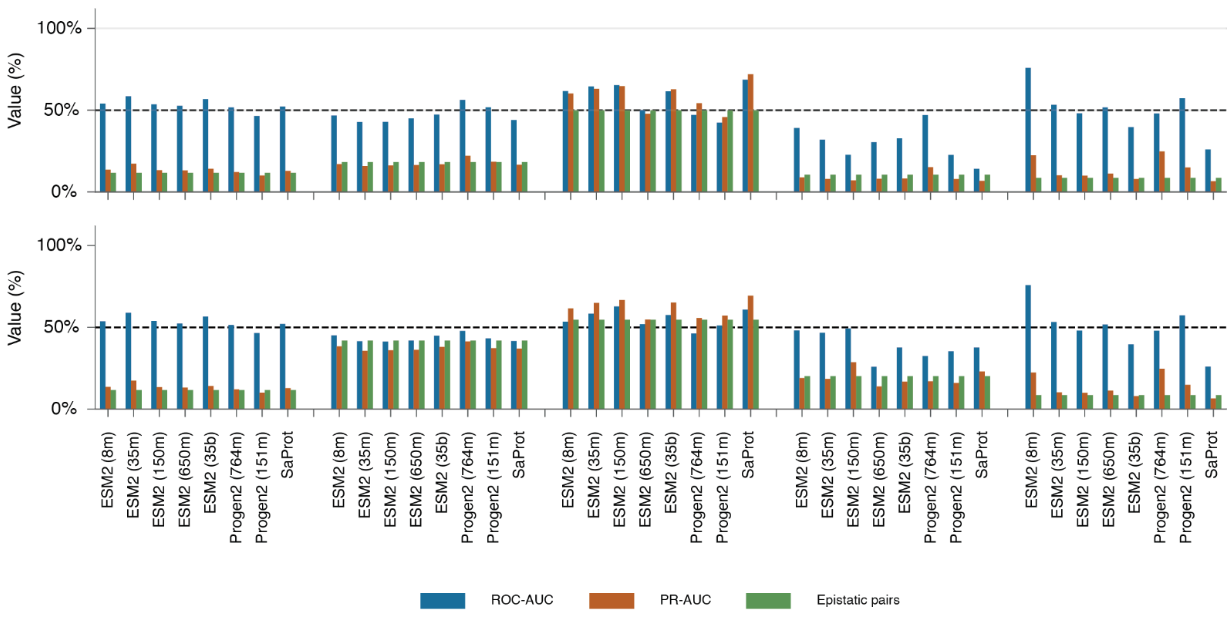
Strong positive + negative epistasis only
Value (%)
100%
50%
0%
ESM2 (8m)
ESM2 (35m)
ESM2 (150m)
ESM2 (650m)
ESM2 (35b)
Progen2 (764m)
Progen2 (151m)
SaProt
ROC-AUC
PR-AUC
Epistatic pairs

**Supplementary Figure 8 - pLM context dependence does not reliably classify variants exhibiting experimentally measured epistasis.**

**(a)** Absolute difference between contextual and fixed-context additive zero-shot scores for experimentally epistatic and non-epistatic mutation pairs. Values were min-max normalized per dataset for visualization purposes only. The top row is restricted to strong positive and negative sign epistasis excluding all pairs in which negative and positive are combined to remove ambiguity, and the bottom row includes all epistasis classes.

**(b)** Ability of the contextual-minus-additive score difference to classify experimentally epistatic pairs. ROC AUC and PR AUC are shown together with the fraction of evaluated pairs that were epistatic. Top, strong positive and negative sign epistasis only excluding pairs in which the singlets are composed from positive and negatives. Bottom, all epistatic pairs. A value of 0.5 is marked for reference for ROC AUC. The LOV and HIS(2) datasets have small number of pairs, and thus ROC AUC values might be spurious, with low PR AUC values

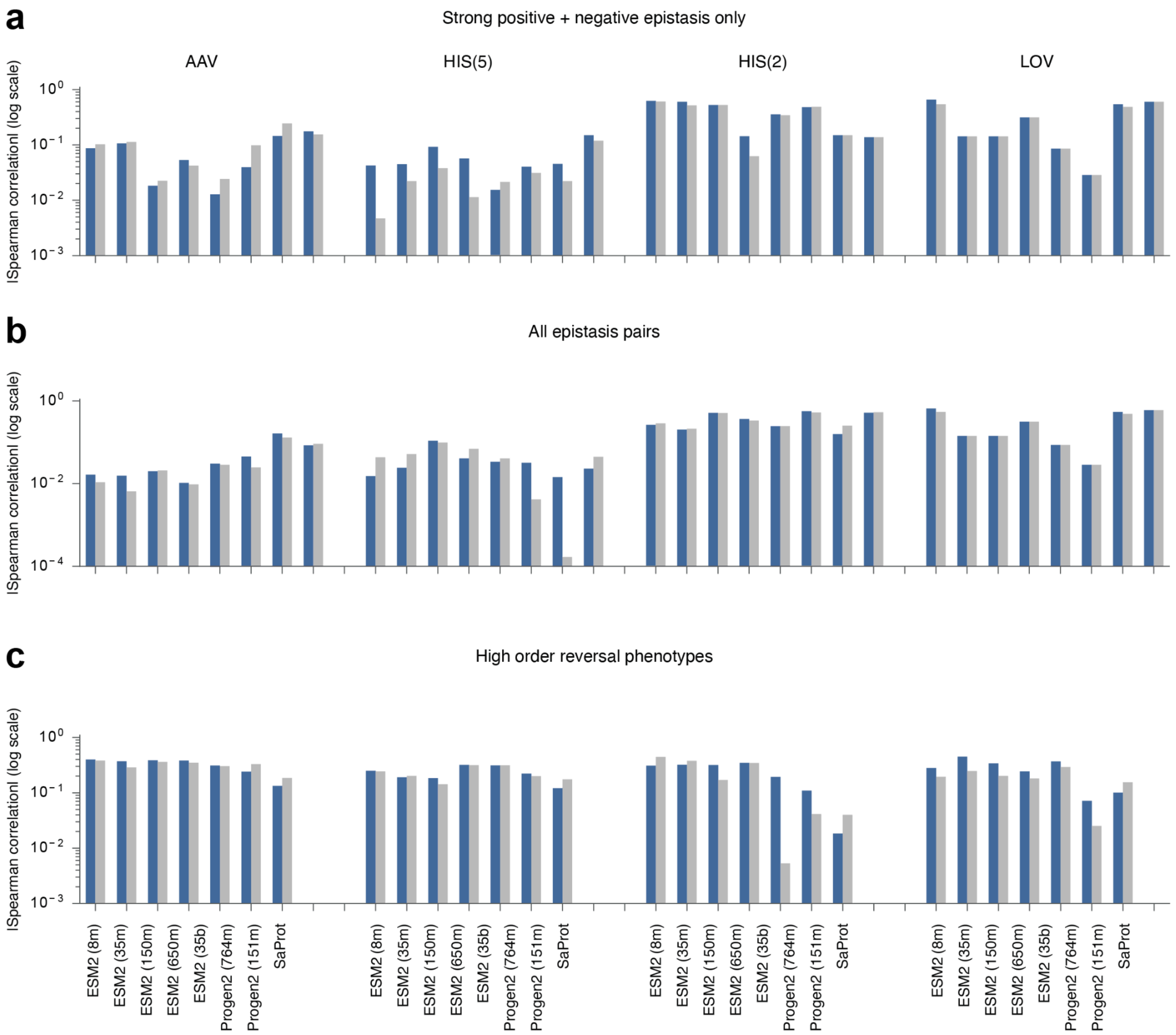
a
Strong positive + negative epistasis only
AAV
HIS(5)
HIS(2)
LOV
|Spearman correlation| (log scale)
b
All epistasis pairs
c
High order reversal phenotypes
ESM2 (8m)
ESM2 (35m)
ESM2 (150m)
ESM2 (650m)
ESM2 (35b)
Progen2 (764m)
Progen2 (151m)
SaProt

**Supplementary Figure 9 - Contextual scoring does not consistently improve phenotype prediction in epistatic variants**

Absolute Spearman correlations between pLM zero-shot scores and experimentally measured phenotypes for variants containing epistatic mutations. Paired bars compare fixed-context additive scoring with contextual, background-dependent scoring across compatible datasets and pLMs.

(a) Variants containing strong positive or negative sign-epistatic pairs, excluding pairs composed of one positive and one negative single mutant.

(b) Variants containing any experimentally identified epistatic pair, including all epistasis classes.

(c) Higher-order variants in which additional mutations reverse the experimentally measured phenotype of an epistatic pair. Correlations are displayed on a logarithmic scale.

Across these groups, contextual scoring does not consistently improve phenotype prediction relative to fixed-context additive scoring.

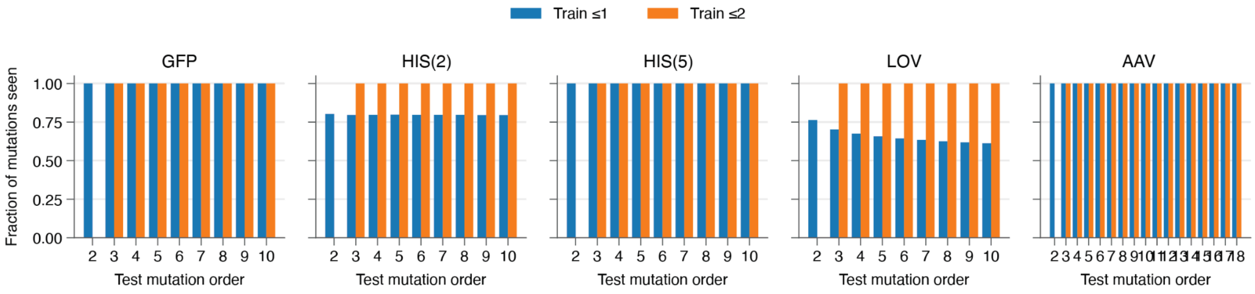


**Supplementary Figure 10 - Coverage of individual substitutions in controlled higher-order extrapolation**

For each held-out test of mutational order, bars show the fraction of mutation occurrences for which the corresponding position and mutant amino-acid identity had already been observed in the lower-order training pool. Blue bars indicate training on measured single-mutant variants (train ≤1), and orange bars indicate training on measured single- and double-mutant variants (train ≤2). Each substitution is counted once per test variant in which it occurs. Panels show GFP, HIS(2), HIS(5), LOV, and AAV.

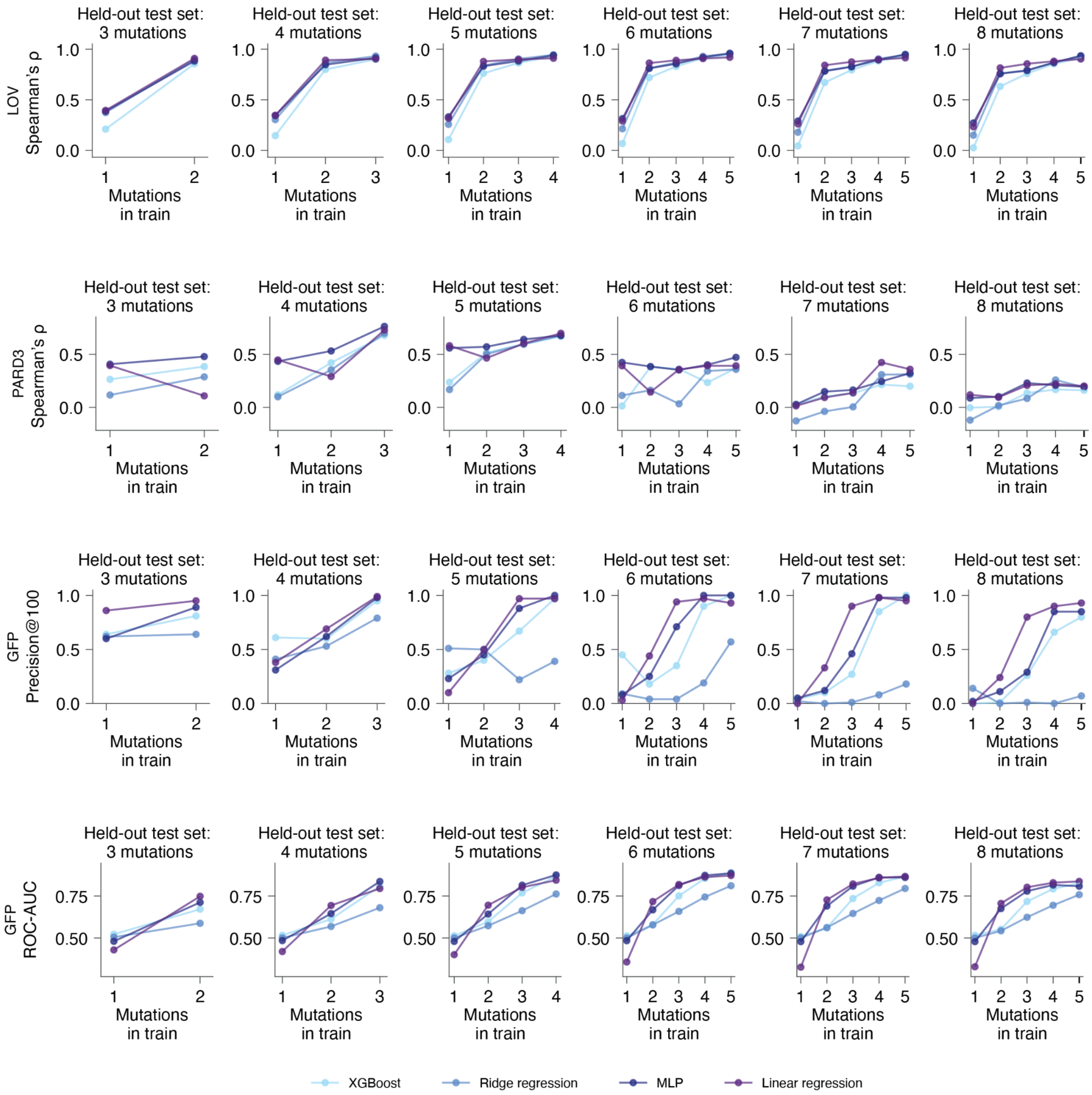

LOV
Spearman's ρ
PARD3
Spearman's ρ
GFP
Precision@100
GFP
ROC-AUC
Held-out test set:
3 mutations
Held-out test set:
4 mutations
Held-out test set:
5 mutations
Held-out test set:
6 mutations
Held-out test set:
7 mutations
Held-out test set:
8 mutations
Mutations
in train
XGBoost
Ridge regression
MLP
Linear regression

**Supplementary Figure 11 - Comparison of supervised prediction algorithms for pLM representations**

Controlled-extrapolation performance of alternative downstream predictors trained on the same pLM-derived representations. Columns correspond to fixed held-out test mutational orders, while the x axis shows the maximum mutational order included in training. Rows show Spearman correlation for LOV and PARD3, Precision@100 for GFP, and ROC AUC for GFP. Linear regression, ridge regression, gradient-boosted trees, and multilayer perceptrons were evaluated on matched train-test partitions. MLPs gave the strongest overall performance and were used for the primary supervised benchmark.

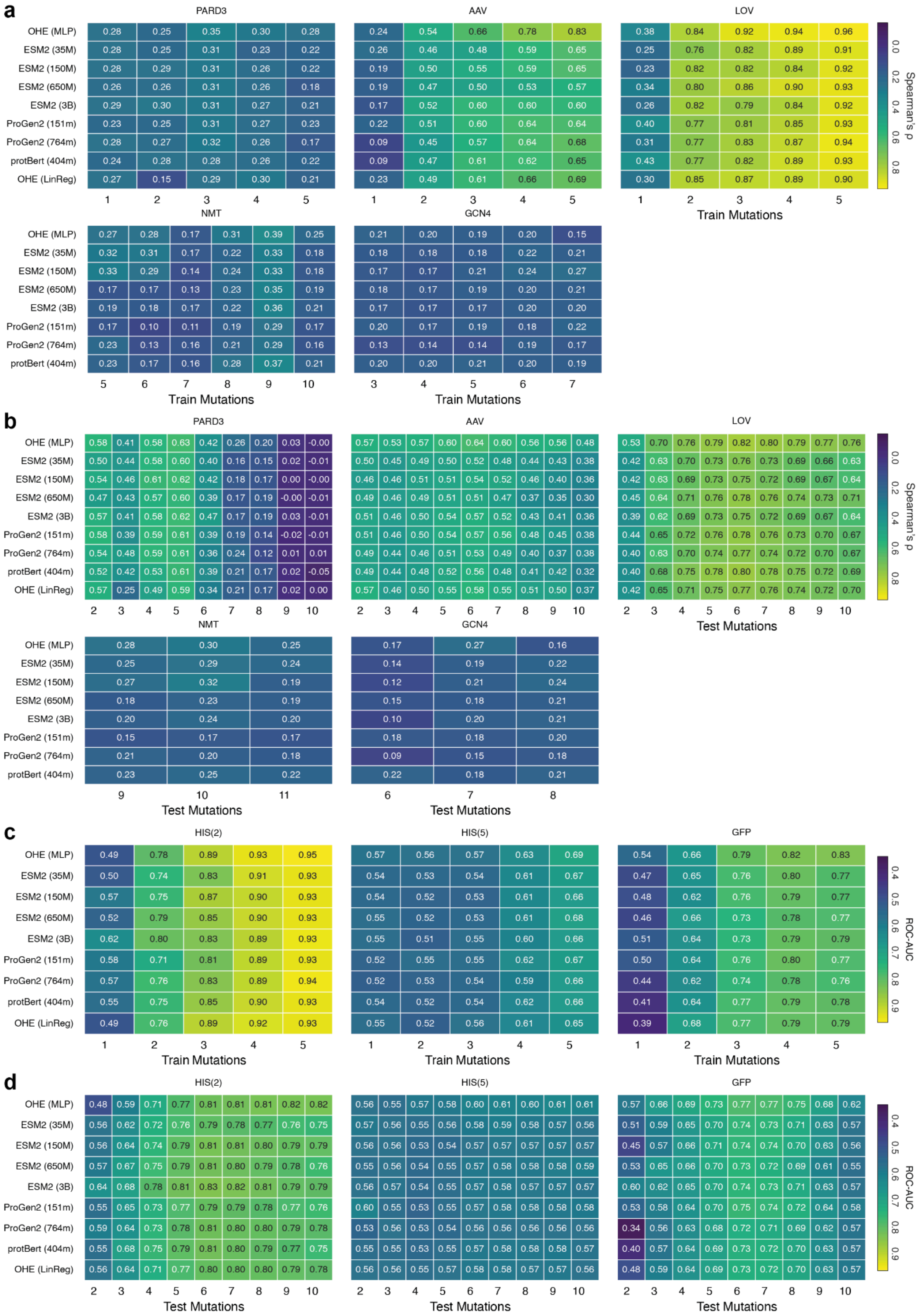
a
b
c
d
PARD3
AAV
LOV
NMT
GCN4
HIS(2)
HIS(5)
GFP
OHE (MLP)
ESM2 (35M)
ESM2 (150M)
ESM2 (650M)
ESM2 (3B)
ProGen2 (151m)
ProGen2 (764m)
protBert (404m)
OHE (LinReg)
Train Mutations
Test Mutations
Spearman's ρ
ROC-AUC

**Supplementary Figure 12 - Controlled-extrapolation performance across models, datasets, and mutational orders**

Heatmaps summarize supervised prediction performance across matched controlled-extrapolation regimes for one-hot encoding (OHE) and protein language model (pLM) representations. Rows correspond to the evaluated representations, including the OHE-based linear regression baseline, and columns indicate the maximum mutational order included in training or the exact mutational order of the held-out test variants. Each cell reports performance averaged across the corresponding eligible train–test regimes.

**(a)** Spearman's ρ for quantitative datasets, grouped by the maximum mutational order included in training. For each training cutoff, performance is averaged across all evaluated higher-order test sets. For example, the OHE (MLP) entry at training order 1 represents the average performance of a model trained on single mutants and evaluated across the available higher-order test sets.

**(b)** Spearman's ρ for the same datasets, grouped by the exact mutational order of the held-out test set. For each test order, performance is averaged across all eligible lower-order training regimes. For example, the ProtBert entry at test order 6 represents the average performance on six-mutant variants when training on variants containing up to one, two, three, four or five mutations.

**(c,d)** Corresponding analyses for the classification datasets GFP, HIS(2) and HIS(5), evaluated using ROC AUC. Panels c and d summarize performance as a function of training and test mutational order, respectively, following the same grouping procedures as in a and b.

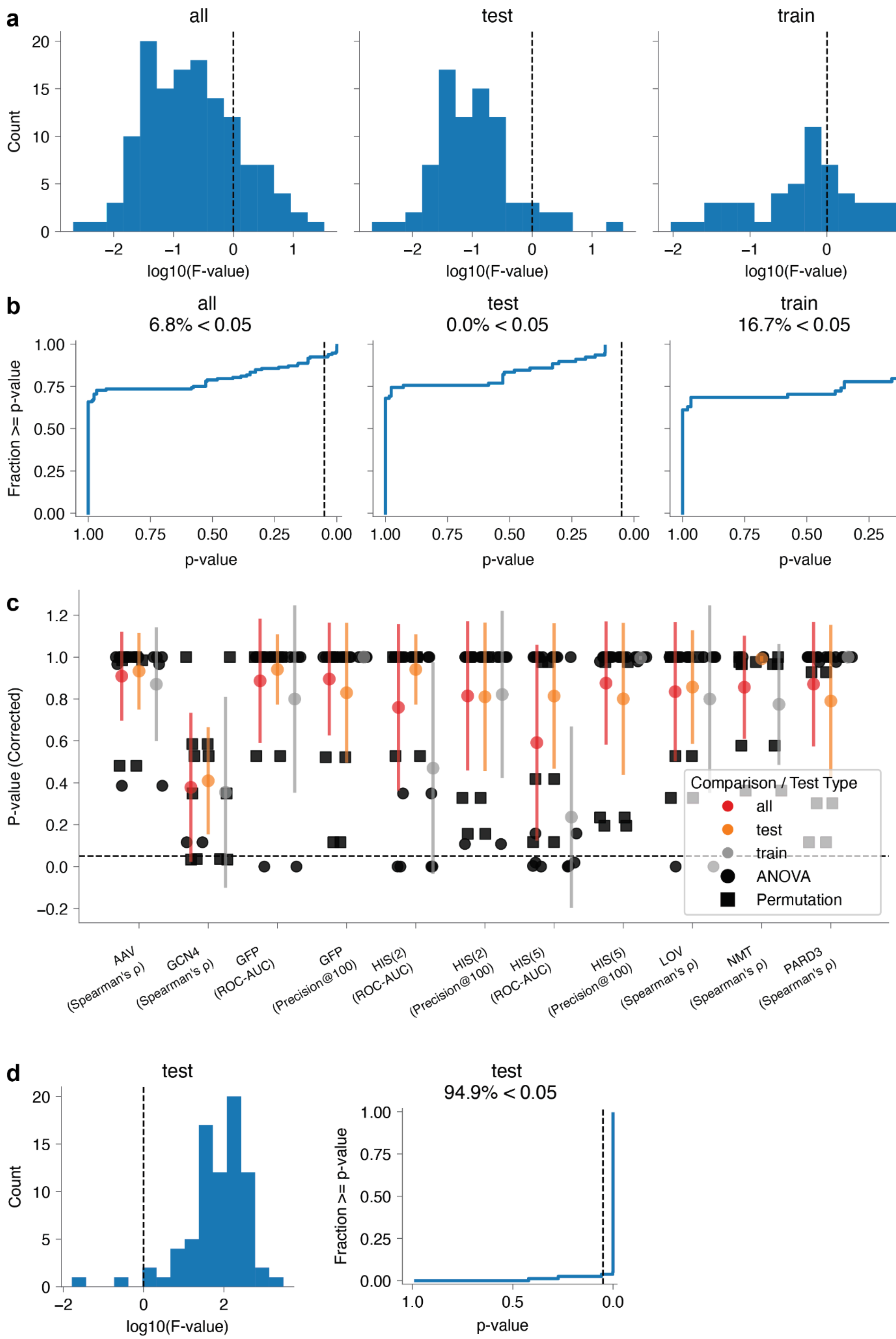
a
all
test
train
Count
log10(F-value)
b
all
6.8% < 0.05
test
0.0% < 0.05
train
16.7% < 0.05
Fraction >= p-value
p-value
c
P-value (Corrected)
Comparison / Test Type
all
test
train
ANOVA
Permutation
AAV (Spearman's ρ)
GCN4 (Spearman's ρ)
GFP (ROC-AUC)
GFP (Precision@100)
HIS(2) (ROC-AUC)
HIS(2) (Precision@100)
HIS(5) (ROC-AUC)
HIS(5) (Precision@100)
LOV (Spearman's ρ)
NMT (Spearman's ρ)
PARD3 (Spearman's ρ)
d
test
Count
log10(F-value)
test
94.9% < 0.05
Fraction >= p-value
p-value

**Supplementary Figure 13 - Statistical comparison of pLM identity and training regimes**

**(a)** Distributions of $\log_{10}$-transformed F-statistics for comparisons of pLM performance across all evaluated conditions (all), fixed held-out test orders (test), and fixed training regimes (train). F-statistics quantify the ratio of between-model to within-model variance. The vertical dashed line indicates $F = 1$.

**(b)** Cumulative distributions of P-values for comparisons of pLM identity across all evaluated conditions (all), fixed held-out test orders (test), and fixed training regimes (train). P-values were adjusted using the Benjamini–Hochberg false discovery rate (FDR) procedure, applied independently to the test-order and training-regime comparison families. The percentages indicate the fraction of significant comparisons (adjusted $P < 0.05$).

**(c)** Dataset-level results for the same comparisons. Each point represents a statistical test of the effect of pLM identity. Colors distinguish comparison schemes, and marker shapes indicate ANOVA or permutation tests. P-values were FDR-corrected as described in (b). The horizontal dashed line marks $P = 0.05$.

**(d)** Corresponding analysis of differences between training regimes evaluated on matched held-out test sets. Left, distribution of $\log_{10}$-transformed F-statistics; right, cumulative distribution of FDR-corrected P-values. Unlike the comparisons between pLMs, differences between training regimes were significant in 94.9% of tests ($P < 0.05$), indicating that the mutational orders represented in training have a substantially greater influence on predictive performance than pLM identity.

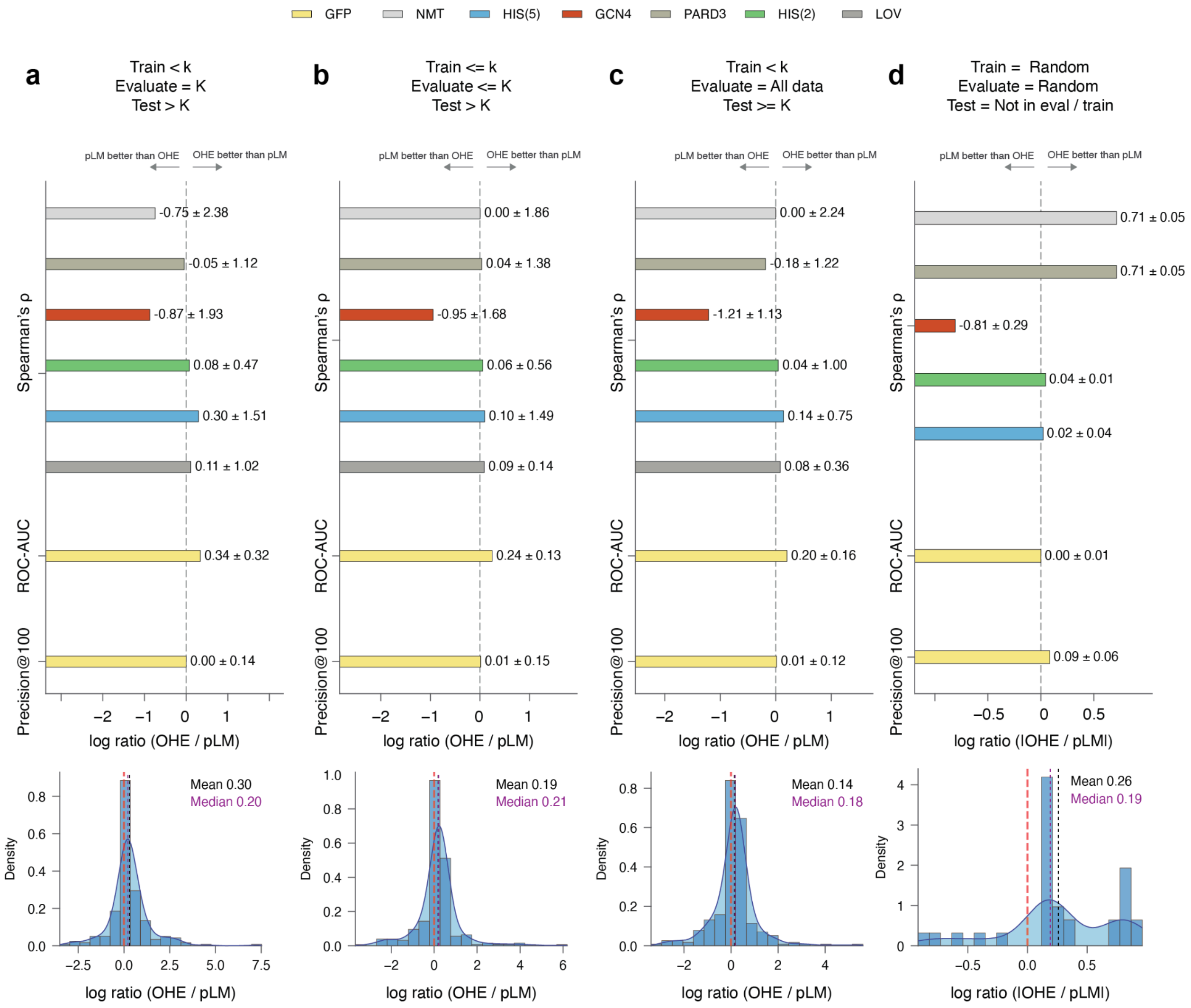

GFP
NMT
HIS(5)
GCN4
PARD3
HIS(2)
LOV
a
Train < k
Evaluate = K
Test > K
b
Train <= k
Evaluate <= K
Test > K
c
Train < k
Evaluate = All data
Test >= K
d
Train = Random
Evaluate = Random
Test = Not in eval / train
pLM better than OHE
OHE better than pLM
Spearman's ρ
ROC-AUC
Precision@100
-0.75 ± 2.38
-0.05 ± 1.12
-0.87 ± 1.93
0.08 ± 0.47
0.30 ± 1.51
0.11 ± 1.02
0.34 ± 0.32
0.00 ± 0.14
0.00 ± 1.86
0.04 ± 1.38
-0.95 ± 1.68
0.06 ± 0.56
0.10 ± 1.49
0.09 ± 0.14
0.24 ± 0.13
0.01 ± 0.15
0.00 ± 2.24
-0.18 ± 1.22
-1.21 ± 1.13
0.04 ± 1.00
0.14 ± 0.75
0.08 ± 0.36
0.20 ± 0.16
0.01 ± 0.12
0.71 ± 0.05
0.71 ± 0.05
-0.81 ± 0.29
0.04 ± 0.01
0.02 ± 0.04
0.00 ± 0.01
0.09 ± 0.06
log ratio (OHE / pLM)
log ratio (|OHE / pLM|)
Density
Mean 0.30
Median 0.20
Mean 0.19
Median 0.21
Mean 0.14
Median 0.18
Mean 0.26
Median 0.19

**Supplementary Figure 14 - OHE remains competitive across alternative model-selection procedures**

Comparison of one-hot encoding (OHE) and pLM-based supervised predictors under alternative training and model-selection procedures. The upper panels summarize the log performance ratio between OHE and pLM predictors across datasets and evaluation metrics, including Spearman's ρ, ROC AUC and Precision@100. Positive values indicate better performance by OHE, whereas negative values favor the pLM predictor. The lower panels show the corresponding distributions of log performance ratios across evaluated train–test regimes, with median values indicated.

**(a)** Extrapolative validation. Candidate architectures were trained on variants containing fewer than K mutations and selected using validation performance on variants containing exactly K mutations. The selected architectures were then evaluated on held-out variants containing more than K mutations.

**(b)** Within-training-regime validation. Training and validation subsets were drawn from variants containing at most K mutations. The selected architectures were subsequently retrained on the available lower-order data and evaluated on held-out variants containing more than K mutations.

**(c)** Optimistic post hoc model selection. Candidate architectures were selected using resampling across the broader experimentally measured landscape. The selected architectures were then refitted using only variants containing at most K mutations and evaluated on higher-order variants. This procedure provides an optimistic sensitivity analysis rather than a prospective model-selection protocol.

**(d)** Conventional random train–test splits. Architectures were selected using internal validation subsets drawn from each randomly sampled training set, without using the held-out test variants. The selected architectures were refitted on the corresponding training sets and evaluated on the held-out test sets.

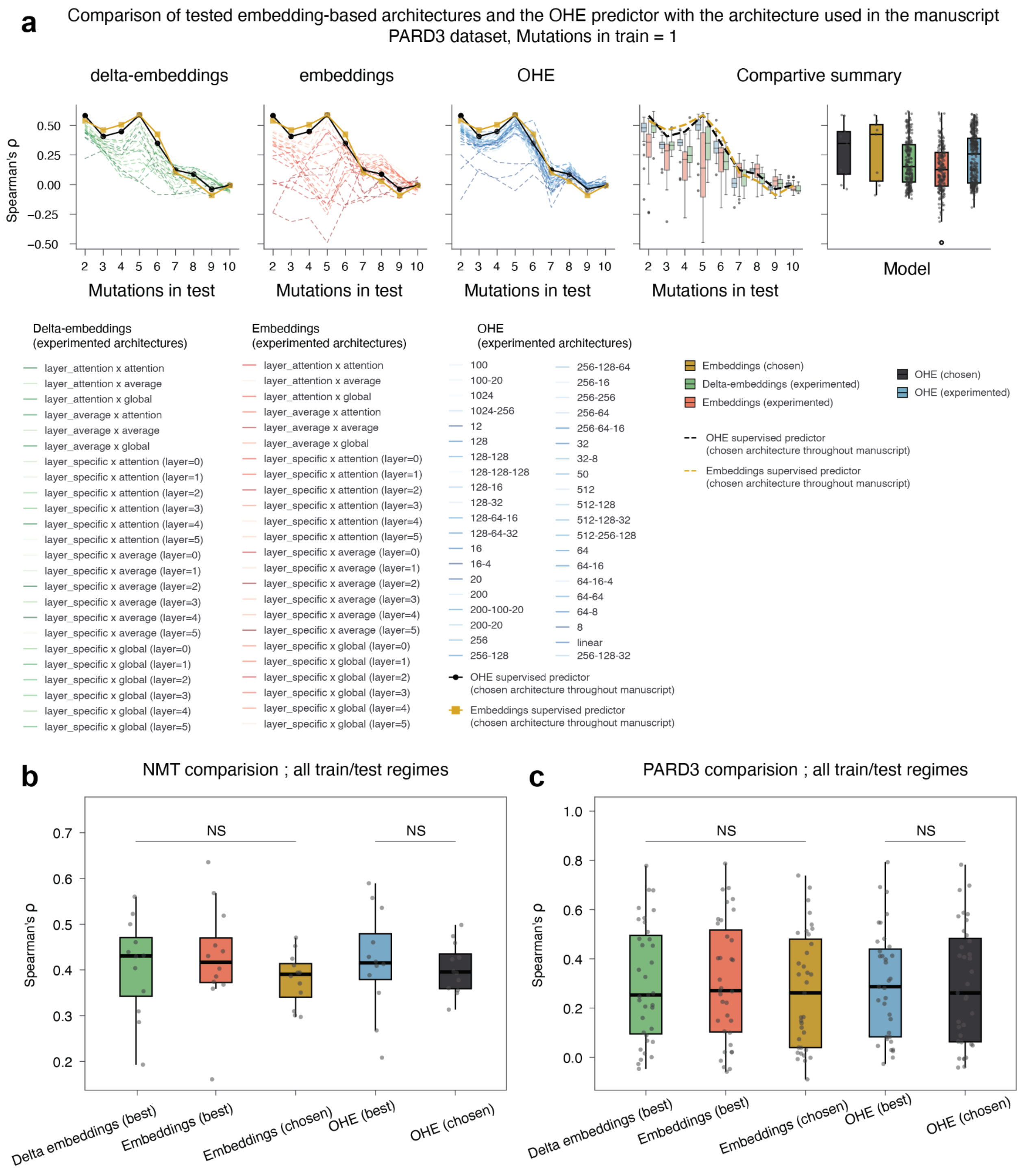
a
Comparison of tested embedding-based architectures and the OHE predictor with the architecture used in the manuscript
PARD3 dataset, Mutations in train = 1
delta-embeddings
embeddings
OHE
Compartive summary
Spearman's ρ
0.50
0.25
0.00
−0.25
−0.50
2 3 4 5 6 7 8 9 10
Mutations in test
Model
Delta-embeddings
(experimented architectures)
layer_attention x attention
layer_attention x average
layer_attention x global
layer_average x attention
layer_average x average
layer_average x global
layer_specific x attention (layer=0)
layer_specific x attention (layer=1)
layer_specific x attention (layer=2)
layer_specific x attention (layer=3)
layer_specific x attention (layer=4)
layer_specific x attention (layer=5)
layer_specific x average (layer=0)
layer_specific x average (layer=1)
layer_specific x average (layer=2)
layer_specific x average (layer=3)
layer_specific x average (layer=4)
layer_specific x average (layer=5)
layer_specific x global (layer=0)
layer_specific x global (layer=1)
layer_specific x global (layer=2)
layer_specific x global (layer=3)
layer_specific x global (layer=4)
layer_specific x global (layer=5)
Embeddings
(experimented architectures)
layer_attention x attention
layer_attention x average
layer_attention x global
layer_average x attention
layer_average x average
layer_average x global
layer_specific x attention (layer=0)
layer_specific x attention (layer=1)
layer_specific x attention (layer=2)
layer_specific x attention (layer=3)
layer_specific x attention (layer=4)
layer_specific x attention (layer=5)
layer_specific x average (layer=0)
layer_specific x average (layer=1)
layer_specific x average (layer=2)
layer_specific x average (layer=3)
layer_specific x average (layer=4)
layer_specific x average (layer=5)
layer_specific x global (layer=0)
layer_specific x global (layer=1)
layer_specific x global (layer=2)
layer_specific x global (layer=3)
layer_specific x global (layer=4)
layer_specific x global (layer=5)
OHE
(experimented architectures)
100
100-20
1024
1024-256
12
128
128-128
128-128-128
128-16
128-32
128-64-16
128-64-32
16
16-4
20
200
200-100-20
200-20
256
256-128
256-128-64
256-16
256-256
256-64
256-64-16
32
32-8
50
512
512-128
512-128-32
512-256-128
64
64-16
64-16-4
64-64
64-8
8
linear
256-128-32
OHE supervised predictor
(chosen architecture throughout manuscript)
Embeddings supervised predictor
(chosen architecture throughout manuscript)
Embeddings (chosen)
Delta-embeddings (experimented)
Embeddings (experimented)
OHE (chosen)
OHE (experimented)
OHE supervised predictor
(chosen architecture throughout manuscript)
Embeddings supervised predictor
(chosen architecture throughout manuscript)
b
NMT comparision ; all train/test regimes
Spearman's ρ
0.7
0.6
0.5
0.4
0.3
0.2
NS
NS
Delta embeddings (best)
Embeddings (best)
Embeddings (chosen)
OHE (best)
OHE (chosen)
c
PARD3 comparision ; all train/test regimes
Spearman's ρ
1.0
0.8
0.6
0.4
0.2
0.0
NS
NS
Delta embeddings (best)
Embeddings (best)
Embeddings (chosen)
OHE (best)
OHE (chosen)

**Supplementary Figure 15 - Supervised results are robust to alternative representation and downstream-model choices**

**(a)** PARD3 controlled-extrapolation analysis comparing alternative pLM-derived representations and downstream architectures with OHE. Tested pLM representations include raw embeddings and delta embeddings, together with alternative transformer-layer and residue-position pooling operations. Thin lines show individual tested configurations and bold traces show the selected configurations used for comparison. The rightmost panels summarize performance across configurations.

**(b,c)** Spearman correlation across all evaluated train-test regimes for NMT and PARD3, respectively. Boxplots compare the best tested delta-embedding representation, the best raw-embedding representation, the representation selected for the main analysis, and the corresponding best and selected OHE predictors. NS denotes a non-significant difference.

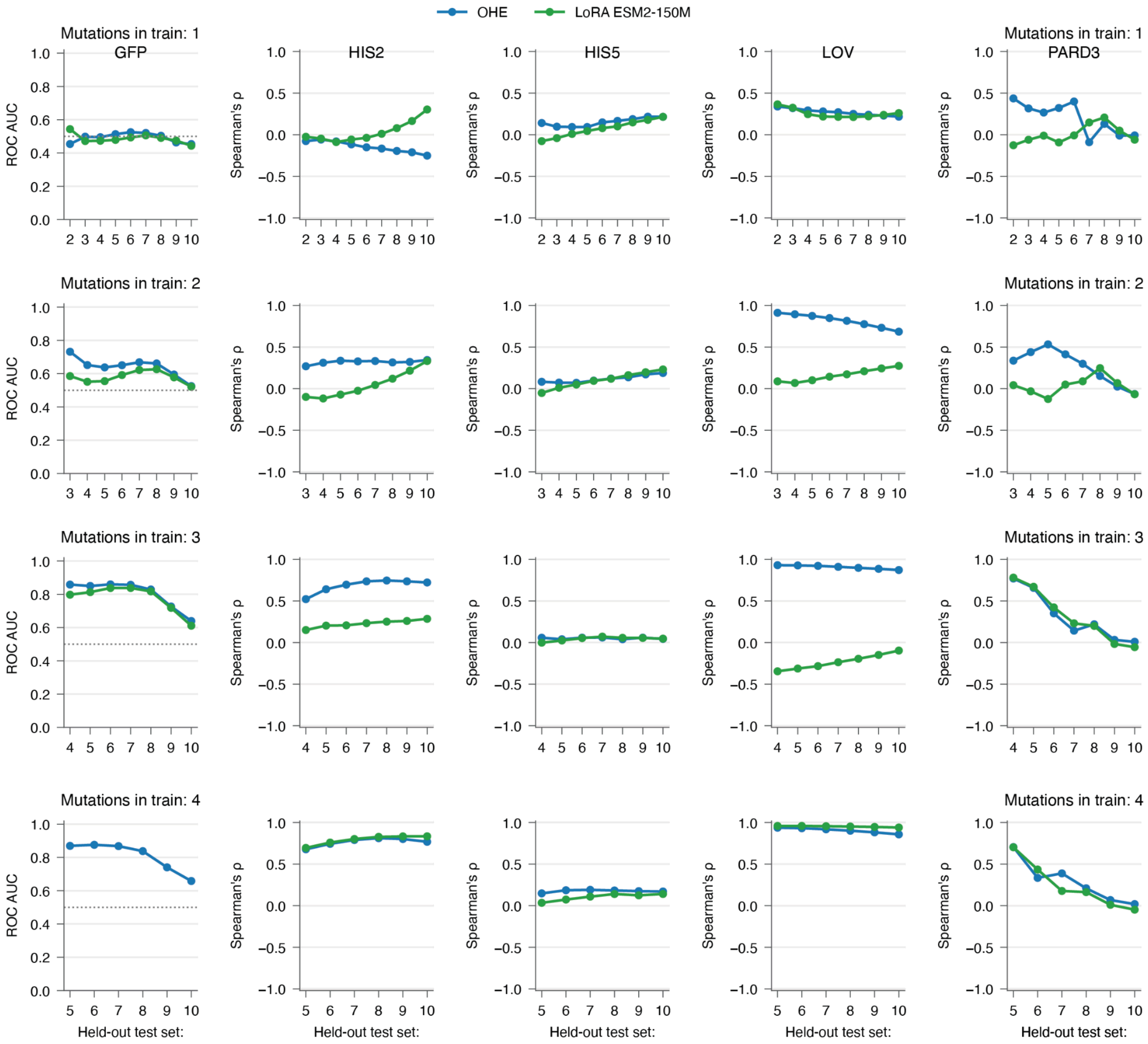


**Supplementary Figure 16 - LoRA fine-tuning under controlled extrapolation across test mutational orders**

Supervised predictors using one-hot encoding (OHE; blue) are compared with ESM2-150M fine-tuned using low-rank adaptation (LoRA; green) across GFP, HIS(2), HIS(5), LOV, and PARD3. Each row corresponds to a fixed maximum training mutational order (1–4), while the horizontal axis shows the mutational order of the held-out test variants. Performance is measured by ROC AUC for GFP and Spearman correlation for the other datasets. For LoRA, validation was performed using variants at the next mutational order relative to the training cutoff.

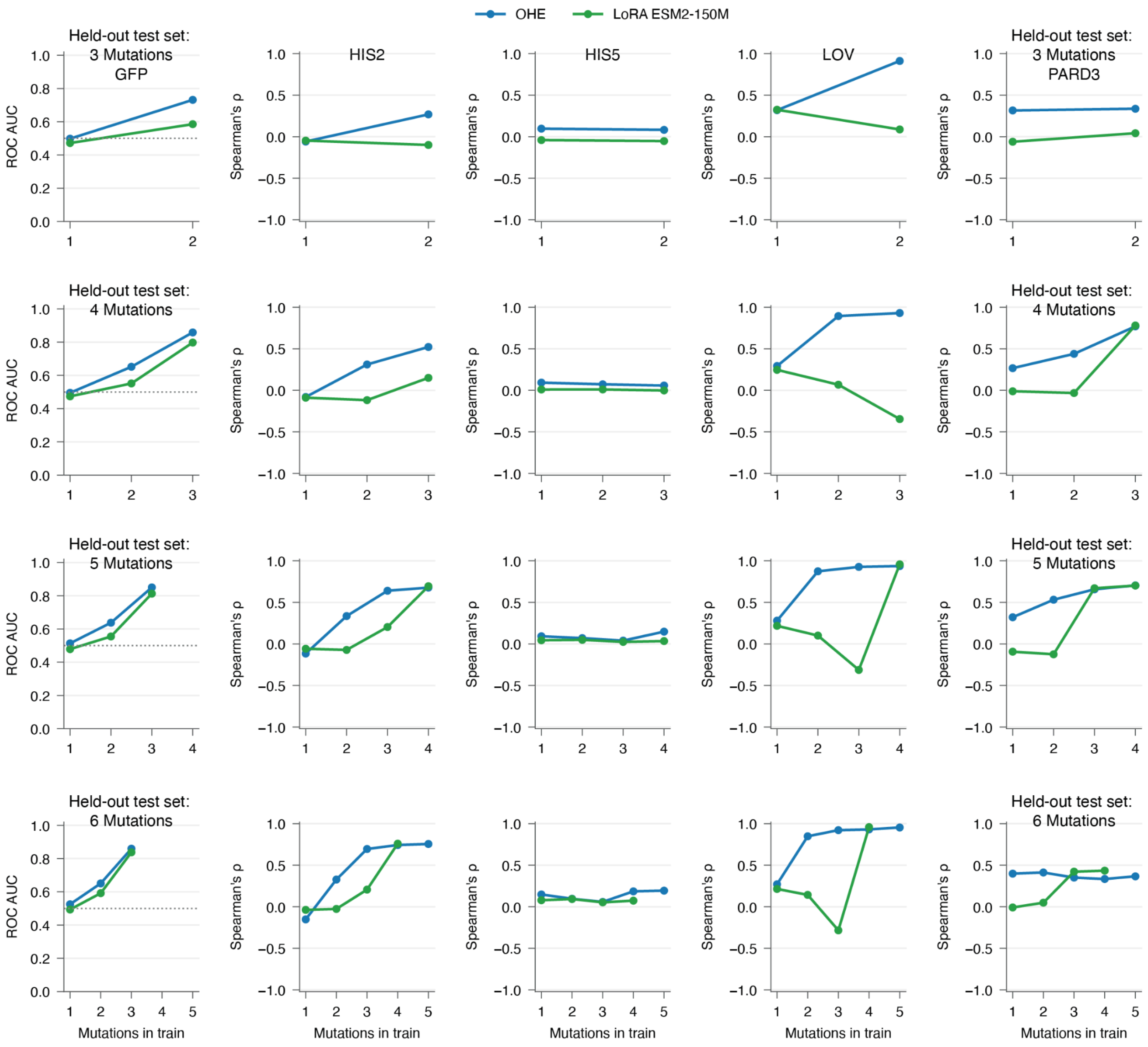


**Supplementary Figure 17 - LoRA fine-tuning under controlled extrapolation across training mutational orders**

Same comparison as in Supplementary Figure 16, but with the held-out test mutational order fixed in each row and the maximum mutational order included in training varied along the horizontal axis. This analysis evaluates how progressively including higher-order variants in training affects prediction of the same held-out mutational order.

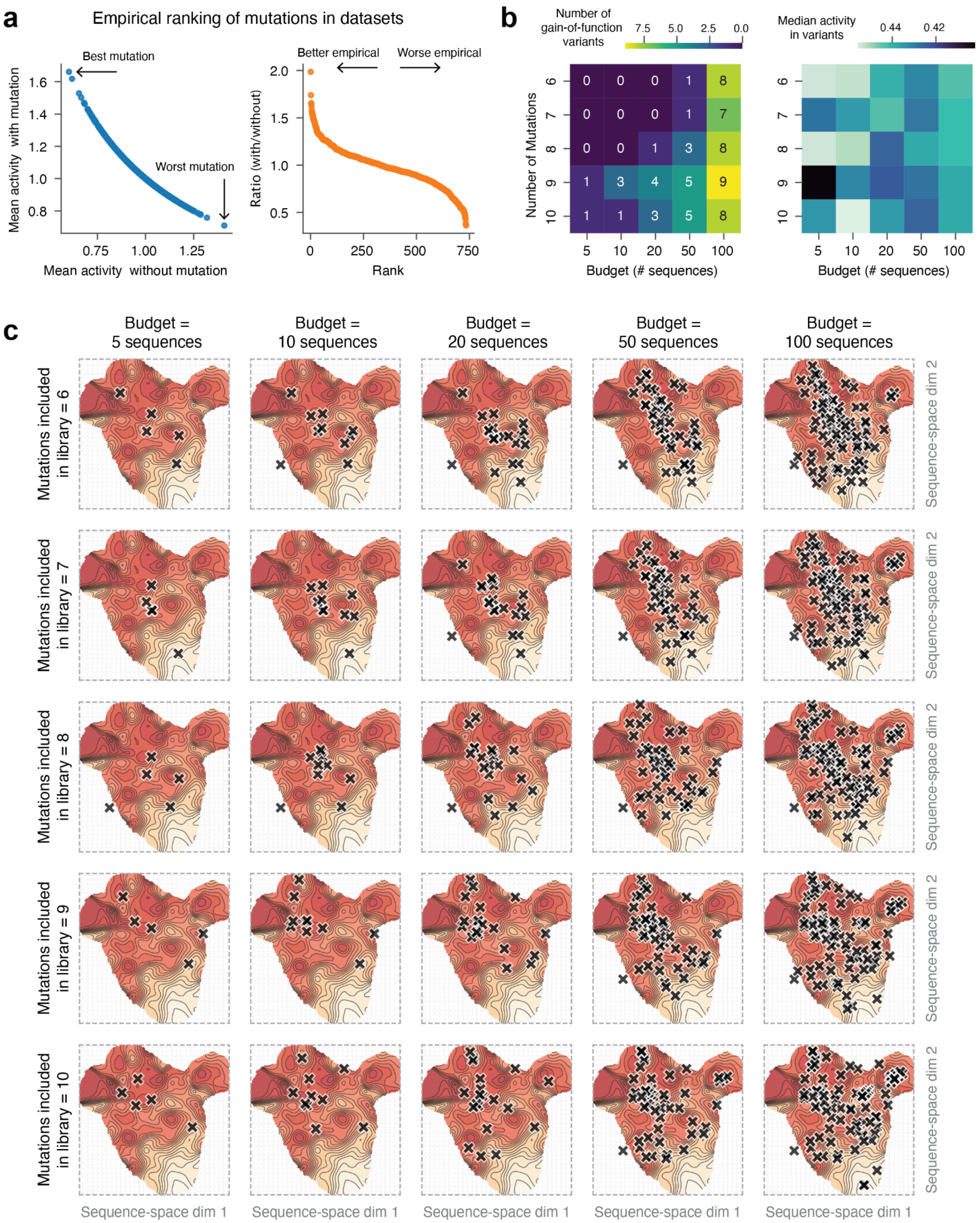
a
Empirical ranking of mutations in datasets
Best mutation
Worst mutation
Mean activity with mutation
Mean activity without mutation
1.6
1.4
1.2
1.0
0.8
0.75
1.00
1.25
Better empirical
Worse empirical
Ratio (with/without)
2.0
1.5
1.0
0.5
0
250
500
750
Rank
b
Number of gain-of-function variants
7.5
5.0
2.5
0.0
Number of Mutations
Budget (# sequences)
6
7
8
9
10
5
10
20
50
100
0 0 0 1 8
0 0 0 1 7
0 0 1 3 8
1 3 4 5 9
1 1 3 5 8
Median activity in variants
0.44
0.42
c
Budget = 5 sequences
Budget = 10 sequences
Budget = 20 sequences
Budget = 50 sequences
Budget = 100 sequences
Mutations included in library = 6
Mutations included in library = 7
Mutations included in library = 8
Mutations included in library = 9
Mutations included in library = 10
Sequence-space dim 1
Sequence-space dim 2

**Supplementary Figure 18 - Definition of the model-guided empirical sub-library screening benchmark**

**(a)** Illustration of empirical mutation ranking. Mutations whose presence is associated with higher average activity than their absence are inferred to occupy more favorable regions of the measured landscape.

**(b)** Example simulated screens. For each screening budget and neighborhood size, different variants are selected from the empirical landscape.

**(c)** Example evaluation of a selected library corresponding to the library shown in (b). Two complementary metrics are reported: the absolute number of gain-of-function variants recovered and the average activity of the selected variants. In the Results section, these quantities are normalized to the optimal library achievable under the same constraints.

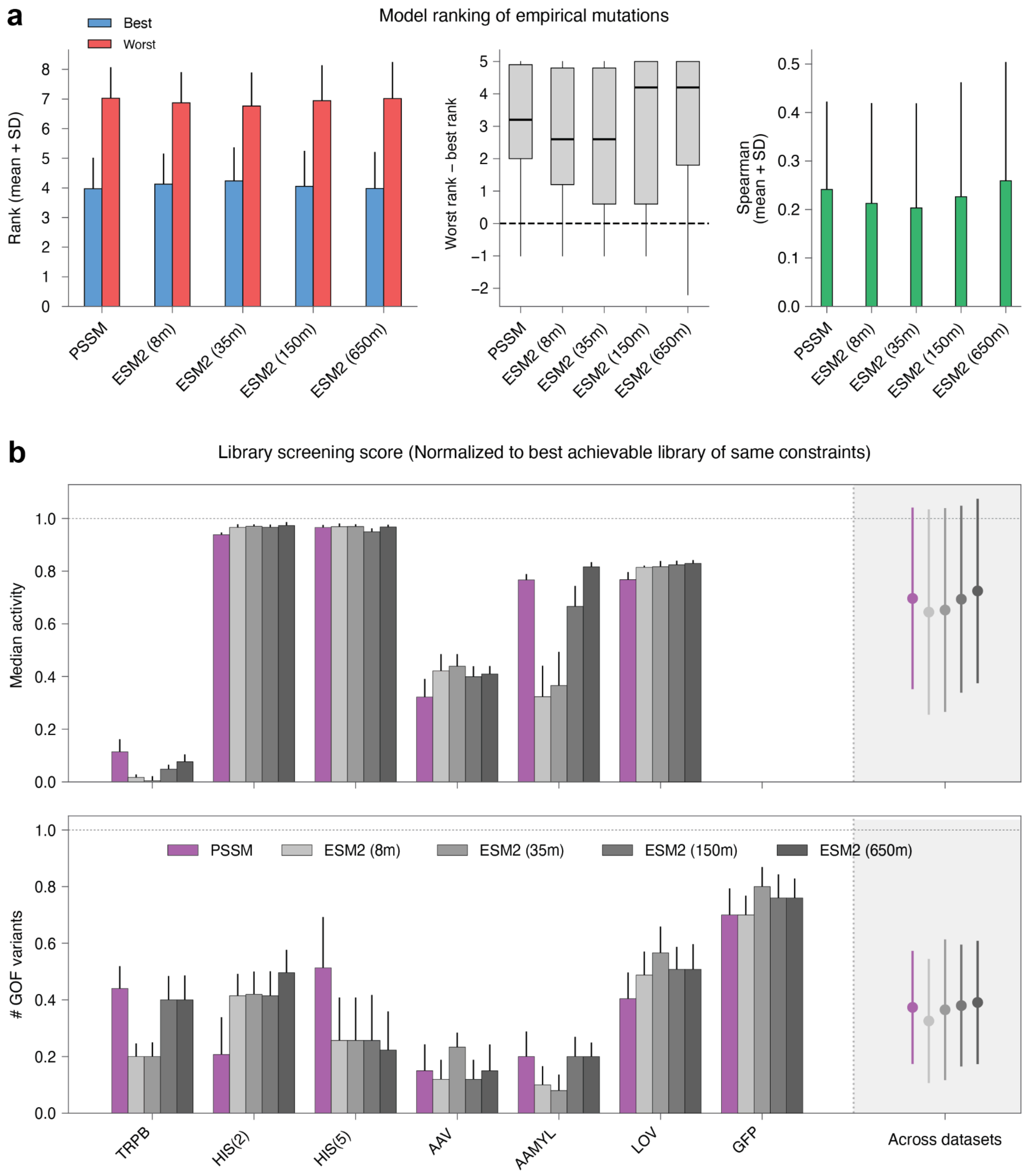

a
Model ranking of empirical mutations
Best
Worst
Rank (mean + SD)
0
1
2
3
4
5
6
7
8
PSSM
ESM2 (8m)
ESM2 (35m)
ESM2 (150m)
ESM2 (650m)
Worst rank – best rank
−2
−1
0
1
2
3
4
5
Spearman
(mean + SD)
0.0
0.1
0.2
0.3
0.4
0.5
b
Library screening score (Normalized to best achievable library of same constraints)
Median activity
0.0
0.2
0.4
0.6
0.8
1.0
### GOF variants
PSSM
ESM2 (8m)
ESM2 (35m)
ESM2 (150m)
ESM2 (650m)
TRPB
HIS(2)
HIS(5)
AAV
AAMYL
LOV
GFP
Across datasets

**Supplementary Figure 19 - Evolutionary and pLM scores provide comparable guidance for mutation ranking and strict sub-library selection**

**(a)** Comparison of model-assigned mutation rankings with experimentally determined mutation benefit. Left, mean ranks assigned to the five empirically most and least beneficial substitutions in each dataset. Middle, difference between the mean ranks of the least and most beneficial substitutions. Right, Spearman correlation between model-assigned and empirical mutation rankings. Empirical mutation benefit was determined from the measured activities of variants containing or lacking each substitution (Methods). Lower model-assigned ranks indicate more favorable predictions. Bars show means ± SD across datasets, with individual datasets indicated by points. Differences between models were not statistically significant (Kruskal–Wallis test; NS, $P > 0.05$).

**(b)** Model-guided sub-library selection under the strict criterion, in which eligible variants contain only mutations from the model-nominated set. Top, median activity of sampled variants. Bottom, number of gain-of-function variants recovered. Performance is normalized to the best empirically identified reference library under the corresponding screening constraints. Bars represent individual datasets, and points on the right summarize performance across datasets.

#### Supplementary Table 1 - Datasets, measured phenotypes and analyses

Abbreviations: ZS, zero-shot ranking; EP, epistasis analysis; CE, controlled extrapolation; RS, random split; EMB, multiple embeddings; LIB, empirical library

| Dataset | Protein | Phenotype / assay | Source / availability | Analyses (original flags) |
|---|---|---|---|---|
| GFP | Green Fluorescent Protein | Fluorescence / original functional activity label | Weinstein et al., 2023; Deposited online | ZS, EP, CE, RS, EMB, LIB |
| PTE | Phosphotriesterase | Phosphotriesterase enzymatic activity | Khersonsky et al., 2018; Deposited online | ZS, RS, LIB |
| NMT | Human nicotinamide N‑methyltransferase | Nicotinamide N-methyltransferase enzymatic activity | Bocquin et al.; provided by authors | ZS, CE, RS, LIB |
| AAV | AAV capsid | AAV capsid packaging fitness | Dallago et al., 2021 (FLIP); public benchmark | ZS, EP, RS, EMB, LIB |
| HIS(2) | HIS3 (region 2) | Growth-based fitness of HIS3 variants | Pokusaeva et al., 2019; ProteinGym | ZS, EP, CE, RS, EMB, LIB |
| HIS(5) | HIS3 (region 5) | Growth-based fitness of HIS3 variants | Pokusaeva et al., 2019; ProteinGym | ZS, EP, CE, RS, EMB, LIB |
| LOV | light-oxygen-voltage | CreiLOV fluorescence | Chen et al., 2023; ProteinGym | ZS, EP, CE, RS, EMB, LIB |
| PARD3 | mesorhizobium opportunistic antitoxin ParD3 | ParD3 antitoxin activity | Ding et al., 2024; ProteinGym | ZS, CE, RS, LIB |
| GCN4 | Transcription Factor GCN4 | Transcriptional activation | Staller et al., 2018; ProteinGym | ZS, CE, RS, LIB |
| TrpB | Tryptophan synthase β-subunit | tryptophan synthase beta subunit enzymatic activity | Almhjell et al., 2024; FLIP2 | LIB |
| AlphaAmyl | α-Amylase | Alpha-amylase enzymatic activity | van der Flier et al., 2024; FLIP2 | LIB |

#### Supplementary Table 2 - Protein language models evaluated in this study

| Model / family | Pretraining objective | Parameters/ hidden dimension | Zero-shot scoring | Training tokens and setup |
|---|---|---|---|---|
| ESM2-8M | Masked LM | 8M / 320 | Yes | 1T tokens ~2 hours on 512 V100 GPUs; 500k steps with 2M tokens batch |
| ESM2-35M | Masked LM | 35M / 480 | Yes | 1T tokens ~8 hours on 512 V100 GPUs; 500k steps with 2M tokens batch |
| ESM2-150M | Masked LM | 150M / 640 | Yes | 1T tokens ~30 hours on 512 V100 GPUs; 500k steps with 2M tokens batch |
| ESM2-650M | Masked LM | 650M / 1280 | Yes | 1T tokens ~150 hours on 512 V100 GPUs; 500k steps with 2M tokens batch |
| ESM2-3B | Masked LM | 3B / 2560 | Yes | 1T tokens 720 hours on 512 V100 GPUs; 500k steps with 2M tokens batch |
| ProGen2-Small | Autoregressive LM | 151M / 1024 | Yes | 180B tokens Direct training details not provided |
| ProGen2-Medium | Autoregressive LM | 764M / 1536 | Yes | 180B tokens Direct training details not provided |
| ProtBert (ProtTrans) | Masked LM | 404M / 1024 | No | 2.5T tokens 230 hours on 512 TPU v3-512; 300K steps with 5M tokens batch + 100K steps 7.8M tokens |
| SaProt-650M | Structure-aware pLM (3Di tokens) | 650M / 1280 | Yes | Not reported in original model table |
